# A Practical Guide to Sparse K-Means Clustering for Studying Molecular Development of the Human Brain

**DOI:** 10.1101/2020.12.31.425014

**Authors:** Justin L. Balsor, Keon Arbabi, Desmond Singh, Rachel Kwan, Jonathan Zaslavsky, Ewalina Jeyanesan, Kathryn M. Murphy

## Abstract

Studying the molecular development of the human brain presents unique challenges for selecting a data analysis approach. The rare and valuable nature of human postmortem brain tissue, especially for developmental studies, means the sample sizes are small (*n*), but the use of high throughput genomic and proteomic methods measure the expression levels for hundreds or thousands of variables (e.g. genes or proteins (*p*)) for each sample. This leads to a data structure that is high dimensional (*p >> n*) and introduces the *curse of dimensionality,* which poses a challenge for traditional statistical approaches. In contrast, high dimensional analyses, especially cluster analyses developed for sparse data, have worked well for analyzing genomic datasets where *p >> n.* Here we explore applying a lasso-based clustering method developed for high dimensional genomic data with small sample sizes. Using protein and gene data from the developing human visual cortex, we compared clustering methods. We identified an application of sparse *K*-means clustering (Robust Sparse *K*-means Clustering (RSKC)) that partitioned samples into age-related clusters that reflect lifespan stages from birth to aging. RSKC adaptively selects a subset of the genes or proteins contributing to partitioning samples into age-related clusters that progress across the lifespan. This approach addresses a problem in current studies that could not identify multiple postnatal clusters. Moreover, clusters encompassed a range of ages like a series of overlapping waves illustrating that chronological- and brain-age have a complex relationship. In addition, a recently developed workflow to create plasticity phenotypes (Balsor et al., 2020) was applied to the clusters and revealed neurobiologically relevant features that identified how the human visual cortex changes across the lifespan. These methods can help address the growing demand for multimodal integration, from molecular machinery to brain imaging signals, to understand the human brain’s development.

## Introduction

As molecular tools have become integrated with human neuroscience, there has been a renewed interest in mapping human brain development. Many studies have compared molecular changes among age groups (Law et al., 2003; Duncan et al., 2010; Pinto et al., 2010; Kang et al., 2011; Siu et al., 2015, 2017; Zhu et al., 2018) using distinct life-span stages that developmentalists have described based on physical, cognitive, and psychosocial maturation (Sigelman and Rider, 2017). However, age-binning assumes that those stages are a good fit for molecular development of the brain. In contrast, other areas of human neuroscience are applying data-driven approaches such as principal component analysis (Bray, 2017) or unsupervised clustering (Lebenberg et al., 2018) to identify age-related changes in brain development. Applying cluster analysis to studying the molecular development of the human brain is challenging because of the limited availability of developmental postmortem tissue samples. Nevertheless, clustering algorithms have been developed for high dimensional biological datasets that have a small sample size (*n*) but measurements from many molecular features (*p*) (e.g. genes or proteins). Here we apply one of those approaches, sparse *K*-means clustering (Witten and Tibshirani, 2010; Kondo et al., 2016), to illustrate a data-driven approach for studying brain development that uses the expression of many genes or proteins to partition samples into age-related clusters. Then we show that clustering can identify aspects of human visual cortex development that are not apparent in typical developmental ontologies.

Cellular and molecular findings from postmortem brain tissue are used as benchmarks for linking age-related changes in non-invasive brain imaging signals with the underlying neurobiology. For example, many imaging studies reference synaptic development measurements (Huttenlocher and Dabholkar, 1997) to account for rapid changes in cerebral cortex MRI signals during the first few years of life. More recently, gene expression databases have been used to identify candidate cellular and molecular features, such as those underlying cortical thinning throughout the life-span (Vidal-Pineiro et al., 2020) or testosterone-related structural properties of the adolescent cerebral cortex (Liao et al., 2021). However, the rare and valuable nature of human postmortem brain samples means that gene expression studies have small sample sizes, especially compared to modern MRI studies that use a population neuroscience approach and aggregate data from hundreds or thousands of subjects (Paus, 2016). The issue of sample size is especially critical for brain development, as even well-established tissue banks (e.g. NIH NeuroBioBank) have fewer than 250 samples for most age groups and fewer than 50 for key ages of child development. Finally, the labour-intensive nature of molecular techniques means that studies can only use a subset of the available samples (e.g. (Pinto et al., 2010) n=28; (Kang et al., 2011) n=57; (Siu et al., 2015, 2017) n=30; (Zhu et al., 2018) n=26). Nevertheless, the high dimensional data collected by molecular studies provide a wealth of information about how the brain changes across the life-span.

Although MRI and postmortem studies of human brain development face different methodological challenges, they share many analytical approaches. Both rely on analyses from the high dimensional toolbox to uncover information relevant to the complexities of brain development. Differences in experimental design, however, place distinct constraints on those analyses. High throughput molecular tools have significantly increased the amount of information obtained from each postmortem sample, generating long lists of gene or protein expression values. Those values represent a vector that describes where each sample exists in a high dimensional space that captures the molecular complexity of human brain development. However, the large number of measurements but small number of samples means that the high dimensional space is sparse with points spread virtually equidistantly across the space. The challenge is to determine how samples cluster together in that sparse space and if those data-driven clusters reflect stages of human development.

Cluster analysis is not new in biology (Eisen et al., 1998; Tamayo et al., 1999; Hastie et al., 2000, 2001), but applying it to postmortem studies of human brain development presents unique problems because of the small sample sizes of those studies. When standard clustering techniques have been used to study gene expression changes in human brain development, clusters are found for regional and prenatal versus postnatal groups, but distinct postnatal clusters matching developmental stages have not been reported (Colantuoni et al., 2011; Kang et al., 2011; Carlyle et al., 2017; Li et al., 2018; Zhu et al., 2018; Disorder et al., 2021). Accordingly, it has been challenging to link cognitive, perceptual, or social-emotional stages and prolonged development found using brain imaging with the underlying maturation of molecular mechanisms in the human brain.

Here we provide a practical guide to sparse clustering that focuses on overcoming the small sample size problem to reveal postnatal patterns of molecular development in the human brain. We introduce sparsity-based clustering, and one approach in particular, sparse *K*-means clustering, developed to address the problem of datasets with a large number of observations from proteins or genes (*p*) but a small number of samples (*n*) resulting in a data structure that is *p>>n* (Witten and Tibshirani, 2010). Finally, we illustrate the value of applying clustering by interrogating the neurobiological features of the clusters to reveal new aspects of the developing human visual cortex.

### Challenges clustering small sample sizes

Currently, transcriptomic, proteomic, and other omics datasets of human brain development include measurements of many molecular features from a small number of samples. The combinatorial nature of those data makes it challenging to use traditional statistical comparisons to understand the many molecular changes that occur in the developing brain. Instead, high dimensional analyses that use all of the data are needed to classify the biological features that differentiate the human brain across the lifespan. However, even when clustering is used, the complexity of the findings can still be challenging to interpret, and studies may need to group the data into predefined age categories to describe the spatiotemporal dynamics of the developing brain (Li et al., 2018).

In the mathematical notation used for clustering algorithms, the genes or proteins are called features or observations and are represented by *p*, while the number of samples is represented by *n*. Most human brain development datasets are either *p ≈ n or p > n* and are best described as high dimensional datasets with more features than samples. When clustering those data, algorithms can *borrow strength* from the large number of features that represent each sample in high dimensional space. However, if only a subset of the features contributes to partitioning the samples into clusters, then the analyses may run into the *curse of dimensionality* (Bellman, 1983). For brain development, this means that developmentally relevant features may become obscured as more and more genes or proteins that do not contribute to developmental changes are included in the dataset. A central problem in analyzing these *p > n* datasets is to identify the molecular features associated with age-related clusters from a very large set of candidate genes. Two approaches for focusing on relevant features include either preprocessing the data using dimension reduction methods (e.g. Principal Component Analysis (PCA), tSNE) or using sparsity-based clustering algorithms that retain all of the features but subset or reweight them during clustering (see Supplementary Material, Box 1. Glossary of Terms).

Some of the common approaches to unsupervised dimension reduction and clustering often used in neuroscience, like PCA and tSNE, can effectively separate data points into clusters in low-dimensional space, especially if there are large differences in features that fall on orthogonal sets of dimensions. For example, tSNE analysis of transcriptomic data identified separate clusters for cortical and cerebellar development (Kang et al., 2011; Carlyle et al., 2017), and PCA has shown that age can explain a large fraction of the variation in protein expression during cortical development (Pinto et al., 2015; Breen et al., 2018). Some of these approaches represent linear combinations of genes or proteins, and focus on reducing dimensionality by identifying correlated features. Problems arise when the features that differentiate clusters are not orthogonal, which may cause linear methods like PCA breakdown and reduce the data onto inappropriate dimensions (Chang, 1983). Thus, traditional dimension reduction and clustering methods are prone to pruning off too much information and, thereby, may miss subtle but significant changes in the human brain’s molecular development. In contrast, sparsity-based clustering methods follow a different approach that keeps all of the features and reweights them in a dissimilarity matrix.

### Approaches to sparsity-based clustering

Because traditional dimension reduction methods may prune off too much information or miss more subtle changes in the human brain’s molecular development, we tested a set of sparsity-based clustering algorithms. Here, sparsity refers to the idea that not all 30,000 genes play a role in brain development and irrelevant dimensions may mask clusters. Furthermore, as more and more features are included, observations become increasingly spread out until they are virtually equidistant. Sparsity-based clustering is a useful approach for analyzing those high dimensional data because the algorithms are not distance-based and can identify a smaller number of molecular features that reflect the spatiotemporal dynamics of neurodevelopment.

In this section, we introduce and compare four clustering methods designed to handle data sparsity but it is not an exhaustive review of sparsity-based clustering.

The agglomerative approach of CLIQUE (Agrawal et al., 1998) finds grids or subspaces in high dimensional data by assigning the desired number of equal length intervals (*xi*) to the grid and a global density value (*tau*) as input parameters. Notably, CLIQUE does not specify the number of clusters in the arguments, but instead compares how many points are in each rectangle of the grid with the overall density parameter and continues to partition the subspaces until the density is less than *tau*. A rectangle in the grid is considered to be dense if the proportion of points in it exceeds the *tau* parameter. CLIQUE then identifies a cluster as the maximal set of dense units in a subspace. For example, using an interval (*xi*) of 2, each dimension of the data is partitioned into 2 non-overlapping rectangles (units) and dense units are identified for further partitioning if they contain a greater proportion of the total number of points than the input value for *tau*. This approach does not strictly partition points into unique clusters and usually results in data points being assigned to more than one cluster. CLIQUE is also prone to classifying points as outliers and excluding them from the analysis.

The divisive clustering of PROCLUS (Aggarwal et al., 1999) is based on medoids and uses a 3-step top-down approach to projected clustering. The steps involve 1) initializing the number of clusters (*k*) and the number of dimensions to consider in the subspace search, 2) iteratively assigning medoids to find the best clusters for the local dimensions, and 3) a final pass to refine the clusters. Typically, PROCLUS has better accuracy than CLIQUE in partitioning points into clusters, but the *a priori* selection of cluster size (*k*) is not easy and demands an iterative approach to finding clusters. Furthermore, by restricting the subspace search size, some essential features may be omitted from the analysis.

Both CLIQUE and PROCLUS were developed for datasets with many more samples (*n*), often 2-3 orders of magnitude larger than most datasets of human brain development. Although those algorithms are accurate for large datasets with thousands of samples, they are less well suited for discovering clusters in small sample sizes. So we needed to test sparsity-based clustering designed for small datasets, and this criteria led us to select two more approaches to sparse hierarchical clustering, SPARCL and RSKC (Kondo, 2016; Witten and Tibshirani, 2018).

SPARCL was developed by Witten & Tibshirani (2010) to adaptively select and reweigh the subset of features during clustering thus eliminating the need for data reduction preprocessing. The algorithm uses a lasso-type penalty to address the challenge of clustering samples that differ on a small number of features. The reweighted variables then become the input to *K*-means hierarchical clustering. The adaptive feature selection of SPARCL focuses on the subset of genes or proteins that underlie differences among clusters, so this process is similar to removing noise from the data. Thus, SPARCL simultaneously clusters the samples and identifies the dominant features thereby making it easier to determine the subset of proteins or genes responsible for partitioning samples into different clusters.

SPARCL has many strengths for analyzing datasets with *p* ≈ *n* or *p > n*; however, it can form clusters containing just one observation (Witten and Tibshirani, 2010). A more recent extension of the algorithm, Robust and Sparse *K*-means Clustering (RSKC), addresses small clusters by assuming that outlier observations cause this problem. RSKC uses the same clustering framework as SPARCL, except that it is *‘robust’* to outliers (Kondo et al., 2016). RSKC iteratively identifies clusters in the data, then identifies clusters with a small number of data points (e.g., n=1) and flags those data points as potential outliers. The outliers are temporarily removed from the analysis, and clustering proceeds as outlined above for SPARCL. Once all clusters have been identified, the outliers are re-inserted in the high-dimensional space and grouped with the nearest neighbour cluster. Thus, RSKC identifies clusters in the data and includes all of the data points.

## Methods

### Datasets

Our lab has been studying development of human visual cortex (V1) by quantifying expression of synaptic and other neural proteins using a library of postmortem tissue samples (n=31, age range 21 days - 79 years, male/female=18/13) (Supplementary Material Table S1). In addition, genome-wide exon-level transcriptomic data that was collected by Kang et al (2011) was used and the postnatal V1 data were extracted (n=48, age range 4 months - 82 years, male/female=27/21) (Supplementary Material Table S2). The transcriptomic data were used to test the reproducibility and scalability of the sparsity-based clustering. The preprocessed exon array data from Kang et al. (2011) were downloaded from the Gene Expression Omnibus (<u>GSE25219</u>). The exon-summarized expression data for 17,656 probes were extracted, and probe identifiers were matched to genes. If a gene was matched by two or more probes and the probes were highly correlated as determined by Kang et. al (2011) (Pearson correlation, r ≥ 0.9), then the expression values were averaged for a total of 17,237 genes.

The clustering methods were tested using three groups of protein or gene data. The first group of protein data was from a series of studies using the Murphy lab postmortem samples to examine the development of molecular mechanisms that regulate experience-dependent plasticity in human V1 (Murphy et al., 2005; Pinto et al., 2010, 2015; Williams et al., 2010; Siu et al., 2015, 2017). Western blotting was run using each sample (2-5 times) to probe for 23 different proteins (Supplementary Material Table S3). The tissue preparation and Western blotting methods have been described in detail previously (Siu et al., 2017, 2018). The initial clustering tests used a subset of 7 proteins (GluN1, GluN2A, GluN2B, GluA2, GABA_A_α1, GABA_A_α3, Synapsin) to explore age-related clustering with AGNES, PROCLUS, CLIQUE, SPARCL and RSKC.

Next, the sparsity-based clustering using RSKC was explored using all 23 proteins to determine how adding more features changed the age-based clustering. Then the reliability of the age-related clustering was explored by running 100 iterations of RSKC with the 23 proteins. A heatmap illustrating the number of times each sample was partitioned into a cluster was made to visualize the reliability.

The scalability of RSKC was tested using a larger protein database and the much larger gene database. These tests included clustering a matrix with 95 proteins collected from the Murphy lab postmortem tissue samples. This time the samples were probed with a high density ELISA array (RayBiotech Quantibody Human Cytokine Array 4000) and an additional 72 proteins were measured for a total of 95 proteins (*p*=95) (Supplementary Material Table S4).

Finally, RSKC clustering was done using the genes in the Kang database by selecting those listed in the SynGO ontology (*n*=988) (Koopmans et al., 2019) and also the full set of genes (*n*=17,237).

### The basic steps to sparsity-based clustering using R

Here we describe four sparsity-based high-dimensional clustering approaches (PROCLUS, CLIQUE, SPARCL, RSKC) for analyzing the development of human V1 using 7 or 23 proteins. Then we explore the scalability of the RSKC method using two larger datasets with 95, 988, or 17,237 proteins or genes.

All of the analyses were done in the R programming language using the integrated development environment RStudio (version 1.3.1093). The basic steps in the workflow used to examine each of the clustering methods are illustrated in Figure 1. The text refers to the R packages that were used and R Markdowns with code and figures are included in the Supplementary Material.

**Figure 1.**
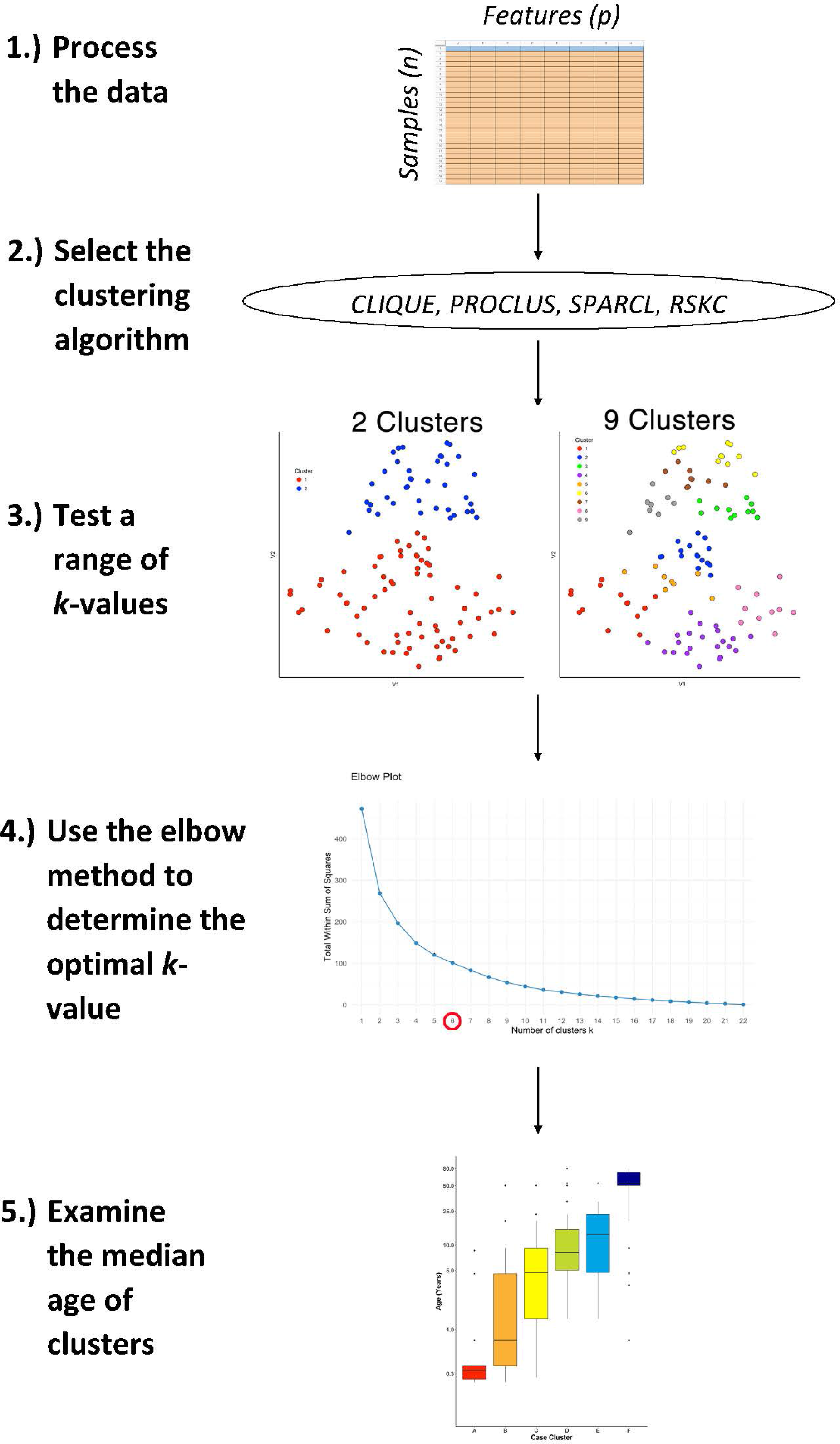
The workflow for studying age-related molecular development of the brain. First, arrange the data into an *n* x *p* matrix, where features (*p*) are represented as columns and samples (*n*) as rows. Then, select the desired sparse clustering algorithm (e.g. CLIQUE, PROCLUS, SPARCL, RSKC) and test its performance along a range of clusters (*k*). Lastly, determine the optimal *k* value using the elbow method and compare the median age of clusters with boxplots.

Figure 1 illustrates the steps that were used for testing various sparsity-based clustering methods to examine if they produce an age-related progression in the median age of clusters. The data were prepared in an *nxp* matrix with each sample forming a row and the features, either proteins or genes, arranged in columns. Those data were used as the input to the clustering algorithms. Here the sparsity-based algorithms tested were PROCLUS and CLIQUE from the *subspace* package (Hassani, 2015), SPARCL (Witten and Tibshirani, 2018), and RSKC (Kondo, 2016). For the algorithms a range of *k* or *xi* values from 2 to 9 were tested to explore the types of clusters produced.

The results of a tSNE dimension reduction was used to visualize clusters for all of the methods tested. However, clustering was not done on the tSNE data itself even though that is a commonly used approach. We used tSNE strictly as a visualization tool because it does a good job of projecting points from high dimensional space onto 2D so that neighboring points reflect their similarity.

The Elbow method was used to determine the number of clusters. Finally, the quality of the age-related clustering of the samples was evaluated by making a boxplot to visualize the progression of the median ages.

This workflow was used for all of the clustering methods described in the next section and an example R Markdown of the analysis is included in Supplementary Material.

## Results

### Evaluating sparsity-based clustering for finding age-related clusters

First, we evaluated the data by exploring if simply visualizing the samples using tSNE produced an age-related organization. The human V1 samples with 7 proteins and all of the WB runs were used as the input to the *tsne* package (Donaldson and Donaldson, 2010) (Fig. 2A). Color-coding the samples by their age showed a global progression in the ages with younger samples mapped to the bottom right and older to the top left in the 2D tSNE space. Next, we applied a commonly used agglomerative hierarchical clustering algorithm, AGNES in the *cluster* package (Maechler, 2019), to test if this clustering approach would reveal age-related groupings of the samples. This algorithm uses the dissimilarity matrix to merge nodes in the tree and it partitioned these data into clusters that suggest an age-related progression (Fig. 2B). However, groups of 2 or 3 adjacent clusters had very similar median ages indicating poor age-related separation of the samples. A major weakness of this hierarchical clustering approach is that incorrect branching can never be undone. Nevertheless, these findings show that even distance-based hierarchical clustering of human V1 postnatal samples can find some age-related progression of postnatal samples.

**Figure 2.**
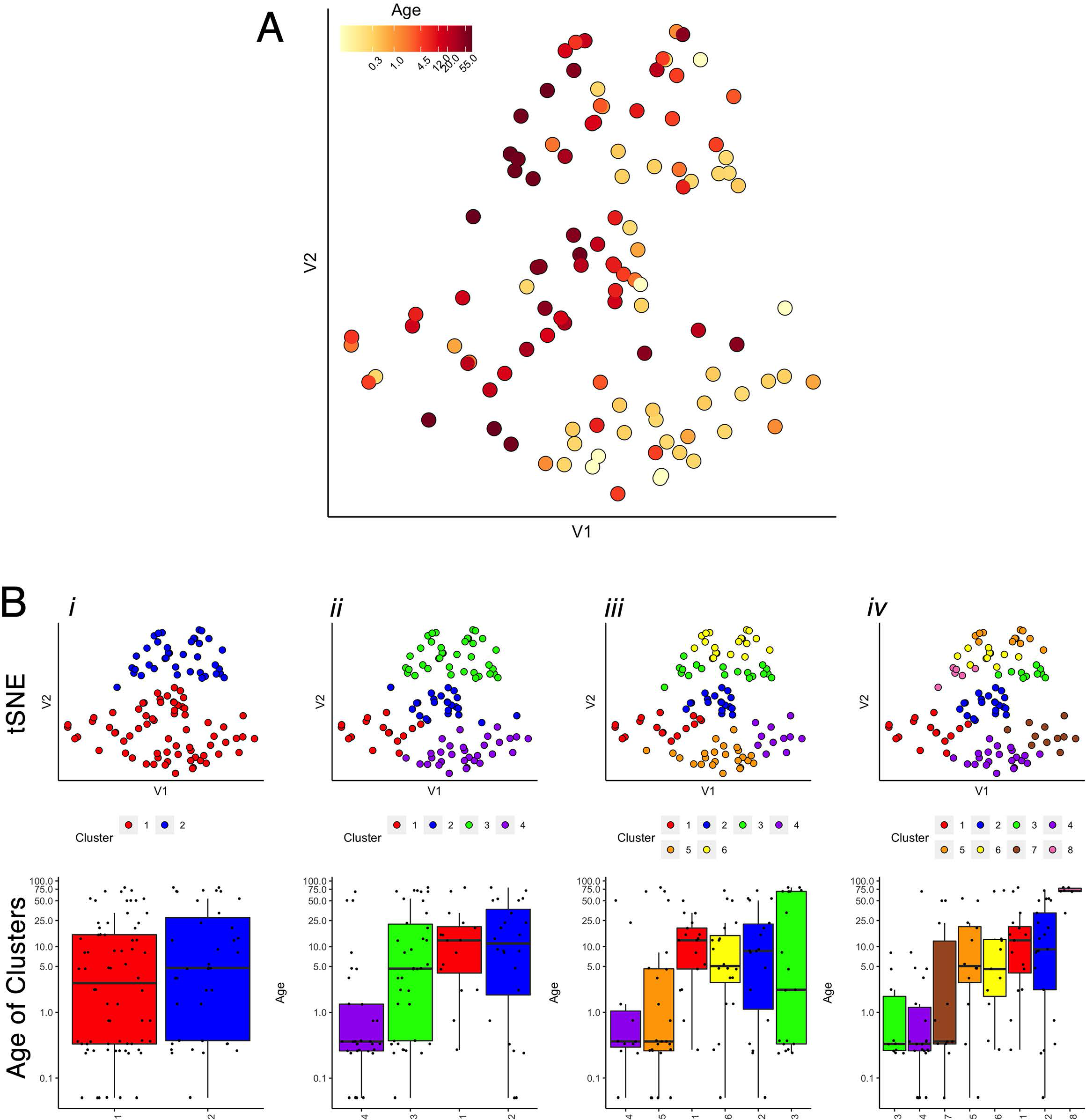
Age-related organization of cases and initial clustering results. A) A 2-Dimensional tSNE scatter plot colour-coded according to individual cases’ age. B) tSNE 2D scatter plots and box plots showing the results of *agnes* for *k* = 2, 4, 6, and 8 case clusters. The tSNE plots display individual samples as points. Points are colour-coded according to their designated cluster determined by *agnes*. Boxplots denote the median and interquartile range of ages in each cluster, and points denote outliers.

Next, we tested the two density projection sparsity-based clustering methods that use either top-down (PROCLUS) or bottom-up (CLIQUE) clustering with all of the observations (n=31) and 7 of the proteins from the human visual cortex development dataset. The outputs were visualized in 2D using tSNE, and the data points were colour-coded according to the clusters identified by each method. Finally, to determine if the clusters represented developmental changes in the dataset, we plotted boxplots showing the median age of the samples in the cluster.

#### PROCLUS

The PROCLUS clustering method was implemented in RStudio using the *ProClus* function in the *subspace* package version 1.0.4 (Hassani, 2015). We explored clusters between *k*=2-9, and Figure 3 shows the results for 2, 4, 6, and 8 clusters for the human V1 data with 7 proteins and all runs included.

**Figure 3.**
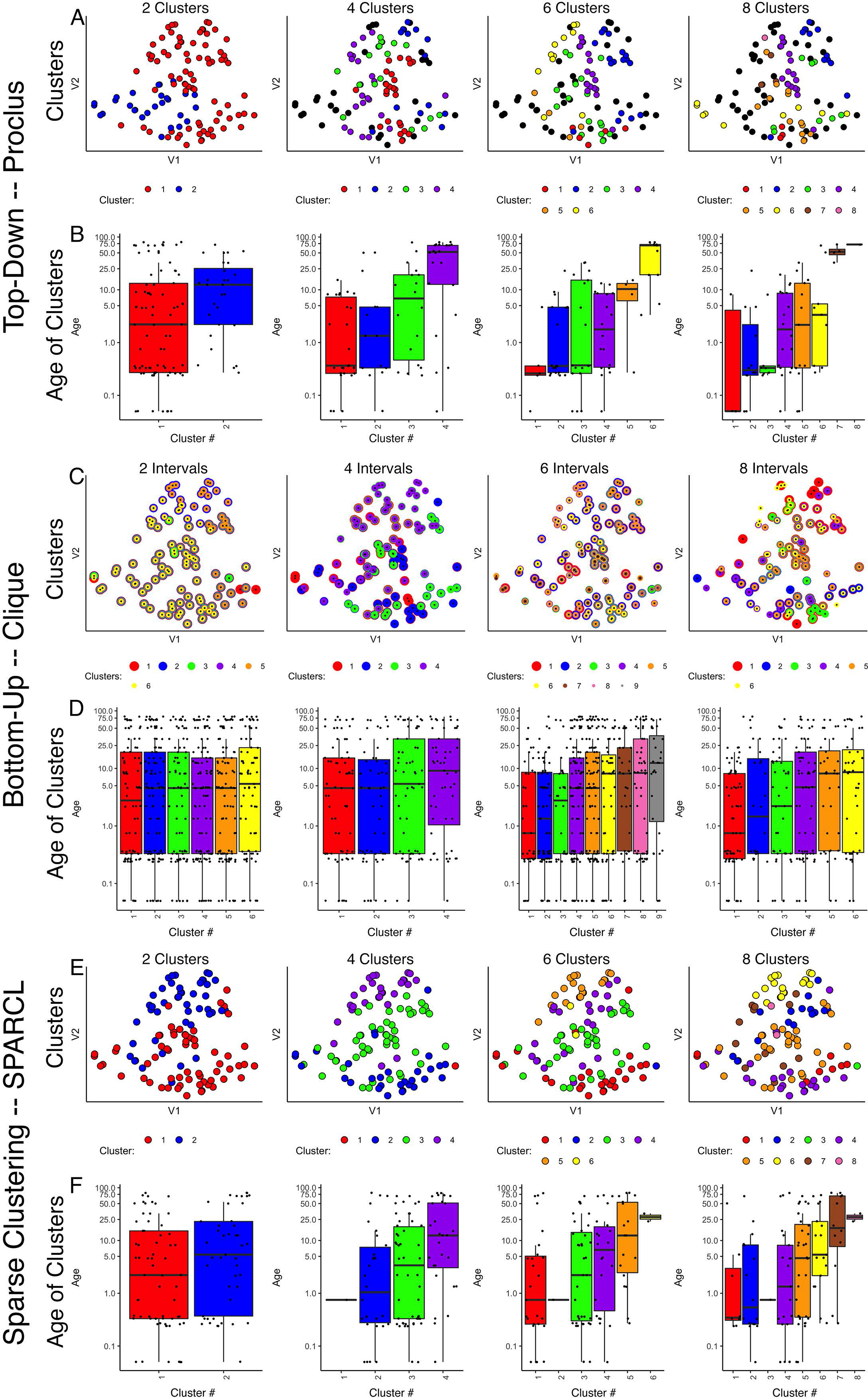
Comparison of various sparse clustering methods. Top-down PROCLUS subspace method across range of cluster numbers (2,4,6,8). The clusters are visualized in tSNE 2D scatter plots of the data by color-coding each data point with its cluster identity (A) and in boxplots showing the median age of the samples in each cluster (B). C-D) Bottom-up Clique subspace clustering method for a range of ‘intervals’. Different clusters are visualized as coloured dots in a tSNE representation of the data (C) and as box plots depicting the mean age of the samples (D). E-F) Sparse clustering after varying the inputted k cluster number (2,4,6,8). Different clusters are visualized as coloured dots in a tSNE representation of the data (E) and as box plots depicting the mean age of the samples (F). The colours in scatter plots and boxplots represent the cluster designation for all plots.

Visualizing the clusters found with PROCLUS (Fig. 3A) showed a mixing of the samples, but the boxplots illustrating the ages of the samples in the clusters suggested an age-related progression, especially for 4 or 6 clusters (Fig. 3B). The PROCLUS clusters’ age progression was somewhat better than the hierarchical clusters but still had clusters with very similar median ages. More importantly, some clusters had only one or two data points, and many samples were tagged as outliers (small grey dots) and excluded from the clusters. Thus, PROCLUS’s iterative top-down feature identification and cluster border adjustments performed poorly for identifying age-related clusters of human V1 development.

#### CLIQUE

The bottom-up clustering method CLIQUE was tested to determine how well this iterative approach to building clusters performed using 7 proteins to group the human V1 samples into age-related clusters.

The CLIQUE function from the *subspace* package (Hassani, 2015) was used to test clustering. CLIQUE requires an input value for the interval setting because the intervals divide each dimension into equal-width bins that are searched for dense regions of data points. Here we tested a range of input interval values (xi=2-8) and those resulted in 4-9 clusters (Fig. 3C&D).

CLIQUE allows data points to be in more than one cluster, so to visualize the multi-cluster identities, we plotted the data points using concentric colour-coded rings. CLIQUE placed all of the data into multiple overlapping clusters, which was true for all interval settings (xi=2-8). The poor partitioning of samples resulted in no progression in the clusters’ median age (Fig. 3D). Thus the iterative bottom-up clustering of CLIQUE performed poorly for clustering the samples into age-related groups.

Comparing these top-down PROCLUS and bottom-up CLIQUE density methods for sparsity clustering showed that neither algorithm was a good fit for producing age-related clustering of the samples. PROCLUS performed somewhat better because some of the parameters resulted in clusters with a progression in the median cluster age; but, the number of data points treated as outliers was unacceptably high.

#### SPARCL

Next, we tested a sparsity-based clustering algorithm, sparse *K*-means clustering, optimized for small sample sizes (Witten and Tibshirani, 2010). The SPARCL package (version 1.0.4) (Witten and Tibshirani, 2018) was used to cluster the human V1 samples with data from 7 proteins. This approach adaptively finds subsets of variables that capture the different dimensions and includes all samples in the clusters. SPARCL searches across multiple dimensions in the data and adjusts each variable’s weight based on the contribution to the clustering. Thus, the term ’sparse’ in this method refers to selecting different subsets of proteins to define each cluster.

To implement sparse *K*-means clustering, we used the *Kmeans.sparsecluster* function in the SPARCL package (Witten and Tibshirani, 2018). We explored a range of *k* clusters between k=2-9. The SPARCL package also includes a function to help determine other input variables, such as the boundaries for reweighting the variables (*wbounds*) to produce optimal clustering.

Visualizing the clusters created by SPARCL showed useful partitioning of the samples into clusters (Fig. 3E) that moved from the bottom right to the top left in the tSNE plot. Also, the boxplots illustrate a good progression of the median cluster age for 4 and 6 clusters. However, SPARCL is prone to making clusters with only 1 sample, and that was the case in this example for *k*=4-9 clusters. To address that problem, we tested another sparse *K*-means cluster algorithm that is robust to making clusters of n=1.

#### RSKC

Finally, we tested a modified version of the SPARCL algorithm called Robust and Sparse *K*-means clustering (RSKC)(Kondo et al., 2016). The RSKC algorithm was designed to be robust to the influence of outliers that can drive other algorithms to create clusters of n=1. RSKC operates by iteratively omitting outliers from cluster analysis, assigning all remaining samples to clusters, and then reinserting outliers to the analysis by grouping them into the nearest-neighbouring cluster.

Using the *RSKC* package in R (Kondo, 2016) we explored clustering for a range of *k* values (*k*=2-9) using the human V1 dataset with 7 proteins and all runs (Fig. 4). The visualization of the clusters on the tSNE plot showed good grouping of the samples into spatially separated clusters. The boxplots illustrate good progression in the median ages of the clusters, especially for 4 or 6 clusters (Fig. 4B). In addition, the algorithm adaptively reweighted the proteins to identify the most robust clusters and we plotted the weights for each of the 7 proteins (Fig. 4C). This component of RSKC identified the lifespan variations in GluN2B, Synapsin, and GluN2A as having the greatest impact on the clustering of the samples.

**Figure 4.**
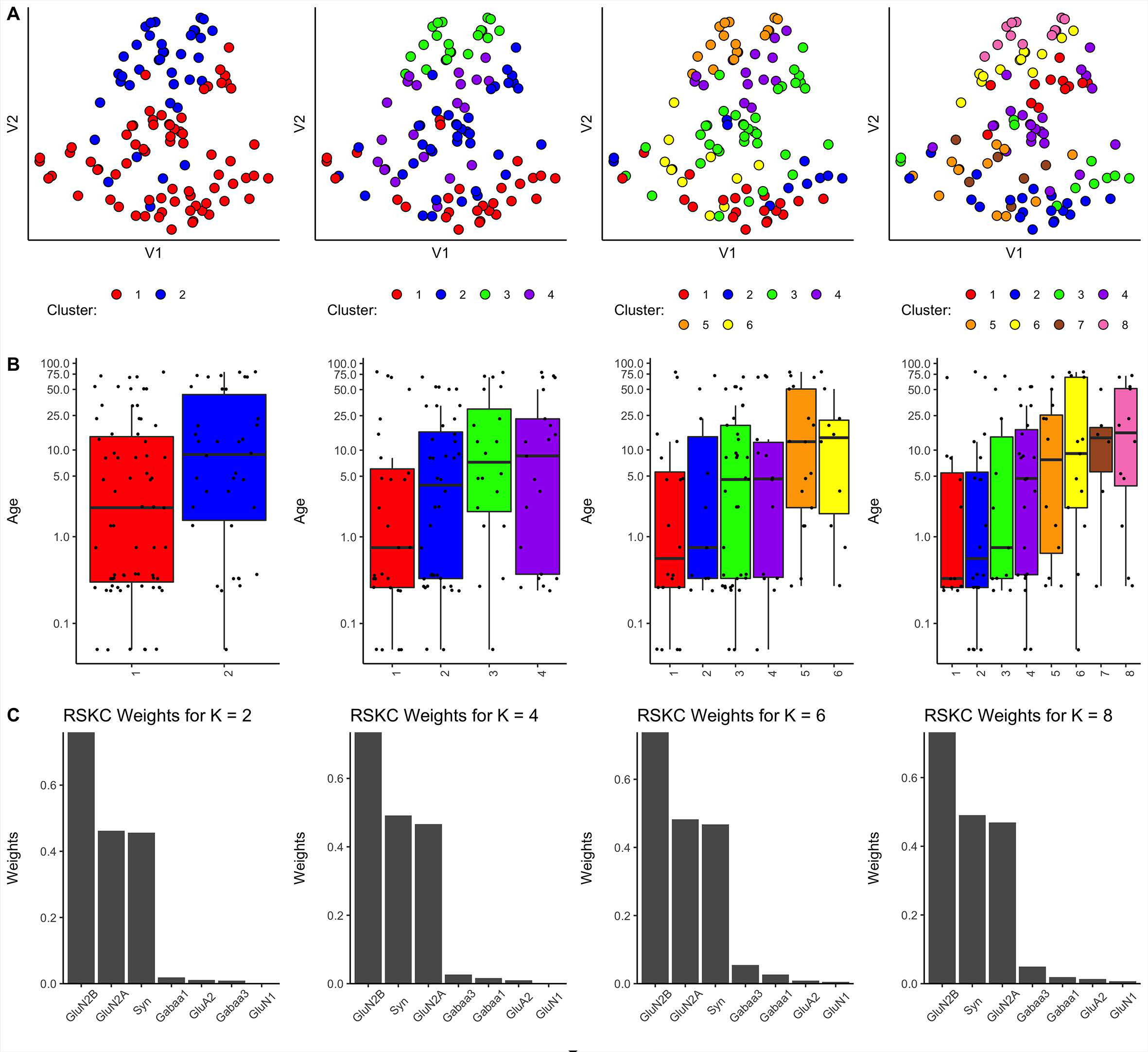
Age-related clustering of 7 synaptic proteins for a single iteration. Expression data from seven synaptic proteins were input into RSKC and used to identify *k =* 2, 4, 6 and 8 case clusters. For each *k*-value, three plots were constructed: A) 2D tSNE scatter plots showing samples colour-coded by their cluster designations, B) Box plots displaying the distribution of ages for each cluster, and C) a bar graph representing the RSKC weights for all seven proteins.

Next, the scalability of RSKC was explored using the full dataset of 23 proteins measured for the human V1 samples (Murphy lab)(Fig. 5). In this example, the average expression value for each protein was used and the elbow plot method identified 6 clusters. Figure 5 shows the results of 3 separate runs of RSKC on the 23 protein dataset. All 3 runs resulted in similar clustering (Fig. 5A-C) with a tight progression of age-related clustering from Cluster A with the youngest median age to Cluster F with the oldest age. The addition of more proteins to the RSKC clustering provided greater precision for identifying the subtle changes that represent the temporal dynamics of human V1 development. The weights for the 23 proteins (Fig. 5D-F) showed that all of the proteins contributed to this high dimensional clustering. Comparing the feature weights among the 3 runs showed some reordering in the weight of individual proteins suggesting that care is needed when using weights from a single run. These weights were used to improve the visualization of the clusters in a tSNE plot. The protein expression values for each sample were transformed by multiplying with the corresponding weight and those transformed data were visualized using tSNE (Fig. 5G-I). Those plots showed the separation of the clusters in the 2D tSNE space.

**Figure 5.**
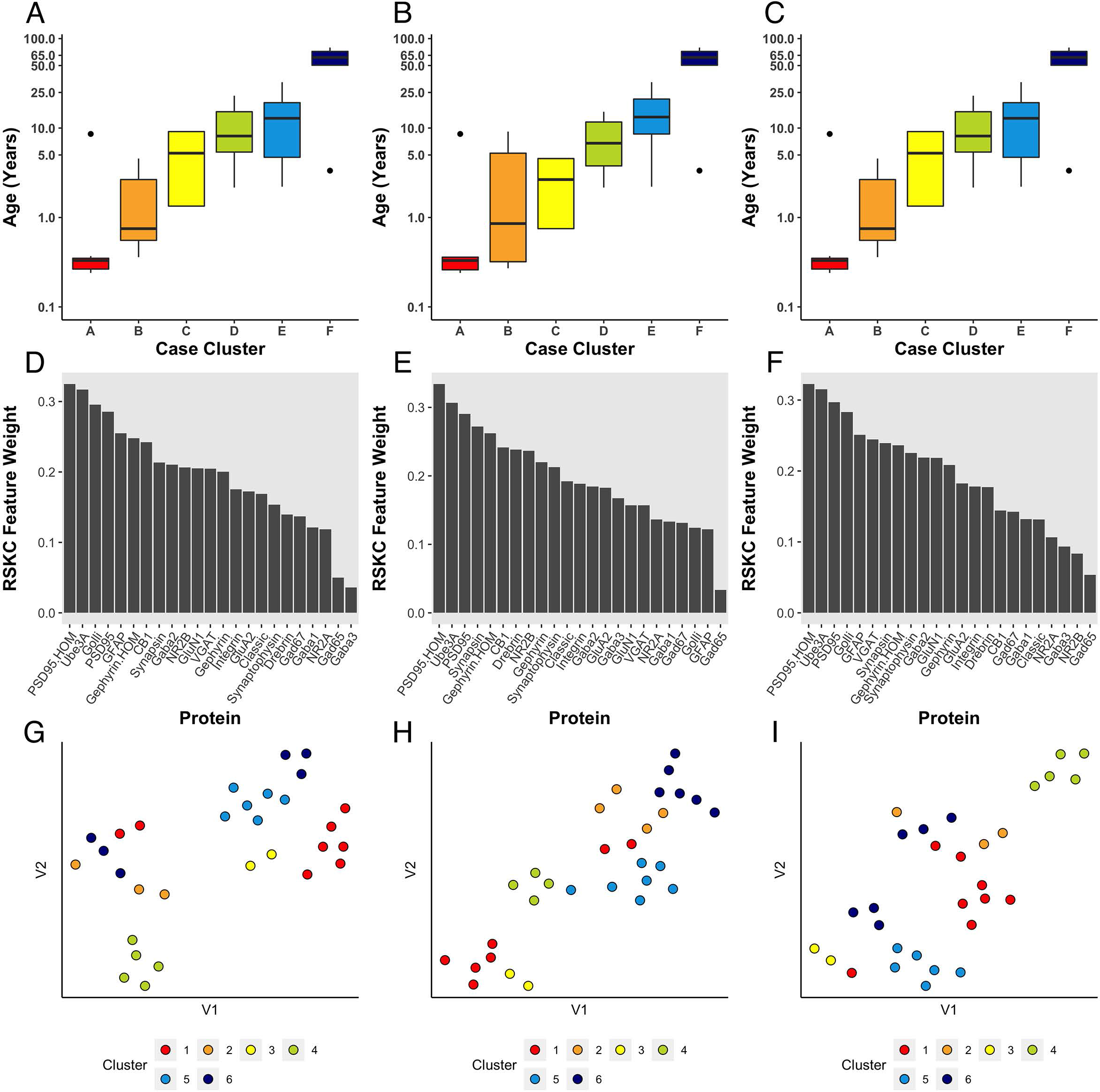
Age-related clustering of 23 synaptic proteins for 3 single iterations. A-C) Expression data from 23 synaptic proteins was used to identify six case clusters. Boxplots of cluster age ordered from youngest (red) to oldest (dark blue) median age. D-F) Bar plot visualizing RSKC feature weights for proteins. G-I) tSNE plot of the protein data scaled by RSKC weights and color-coded by RSKC cluster. In both A-C) and G-I), sample ages were reduced to sample averages to reduce crowding.

Since the starting conditions for clustering can affect which samples end up in a cluster, we tested how robust RSKC clusters were by running the algorithm 100 times with different starting conditions. We then plotted the results of 100 iterations in a boxplot showing the age-related clusters and a heatmap showing the number of times each sample fell into the different clusters (Fig. 6A & B). This analysis showed that the progression in the age of the clusters was robust to the starting conditions (Supplementary Table S5). Furthermore, the heatmap showed that clusters B and C were the least stable, but the other clusters had strong consistency for which samples were partitioned into those clusters. The Jaccard similarity was calculated for all cluster pairs to determine the proportion of samples shared between the clusters. Cases were counted as shared when the cases were partitioned to the cluster 10 or more times because the metric is sensitive to small samples sizes. The similarity indices ranged from 0% to 22% (adjacent pairs: A-B 12%, B-C 22%, C-D 11%, D-E 20%), with cluster C having the most cases shared with other clusters. In addition, the average feature weight for each of the 23 proteins was calculated from the 100 runs (Fig. 6C) and illustrated the gradual progression of feature weights.

**Figure 6.**
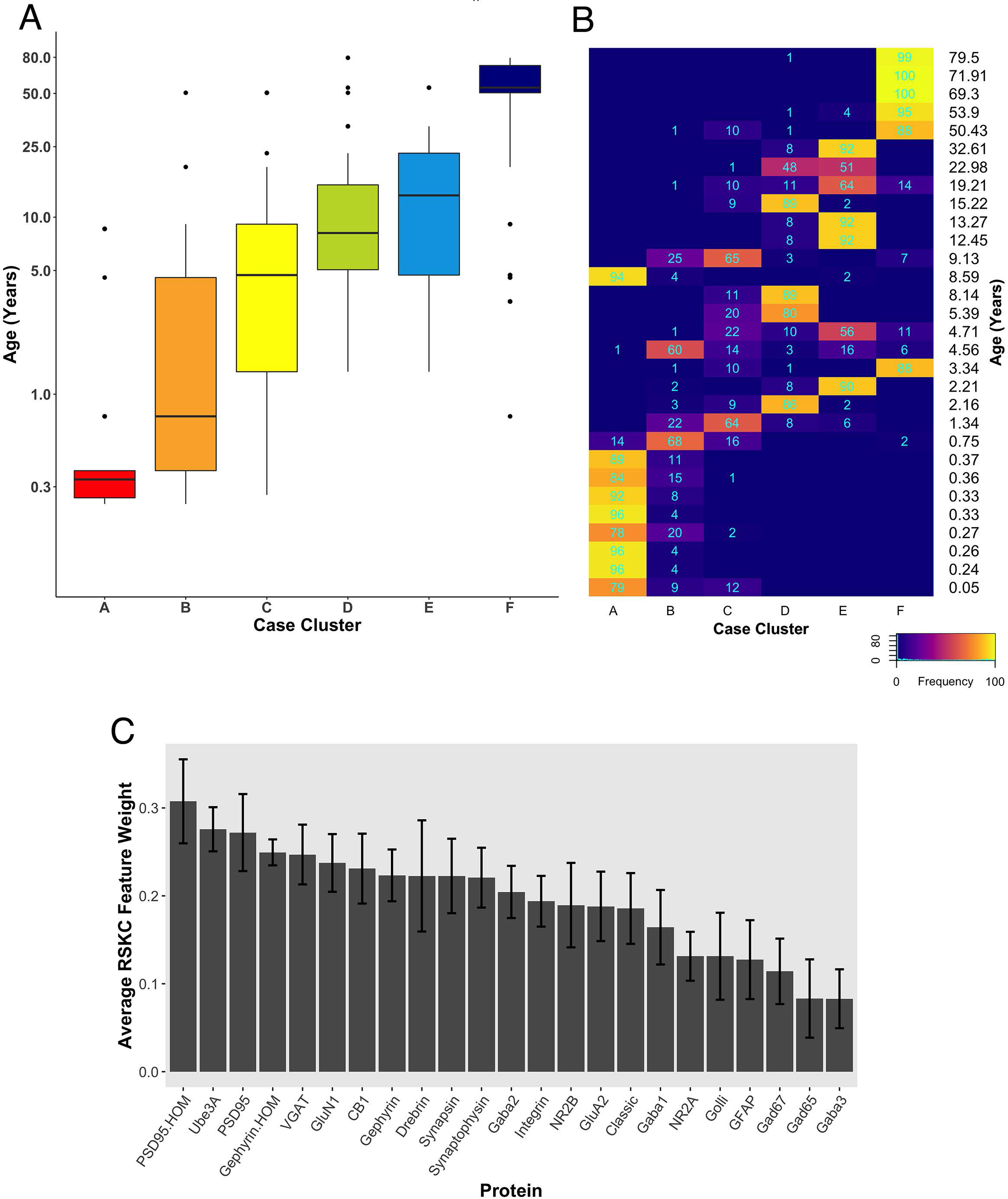
Robust, age-related clustering of 23 synaptic proteins for 100 iterations. A) Expression data from 23 synaptic proteins was used to identify six case clusters. The cluster designation of each case over 100 iterations of RSKC was used to visualize the distribution of case ages (in years). Boxplots denote the median and interquartile range of ages in each cluster, and points denote outliers. B) Heatmap visualizing the number of times each case was assigned to each cluster over 100 iterations of RSKC. C) The average RSKC feature weight for each of the proteins from the 100 iterations.

### Testing RSKC with larger numbers of proteins or genes

So far, we have shown that RSKC does a good job of partitioning samples into age-related clusters with datasets that have fewer than 25 proteins. Here we examine if RSKC scales to larger datasets with 2 to 3 orders of magnitude more features.

We ran the RSKC clustering using data collected from the Murphy lab human V1 samples with measurements for 95 proteins (Supplementary Material Table S4)(Fig. 7A). Once again, 100 iterations of RSKC clustering was used to ensure that the clusters were robust to the starting condition. This analysis found strong age-related clustering of the samples showing 6 well-defined clusters that stepped across the lifespan. We tested if the progression of cluster ages could arise by chance by rerunning the clustering but on each iteration the age of the sample was randomized. As expected, randomizing the ages resulted in clusters with a very broad range of ages and no progression in the mean cluster age (Supplementary Fig. S1).

**Figure 7.**
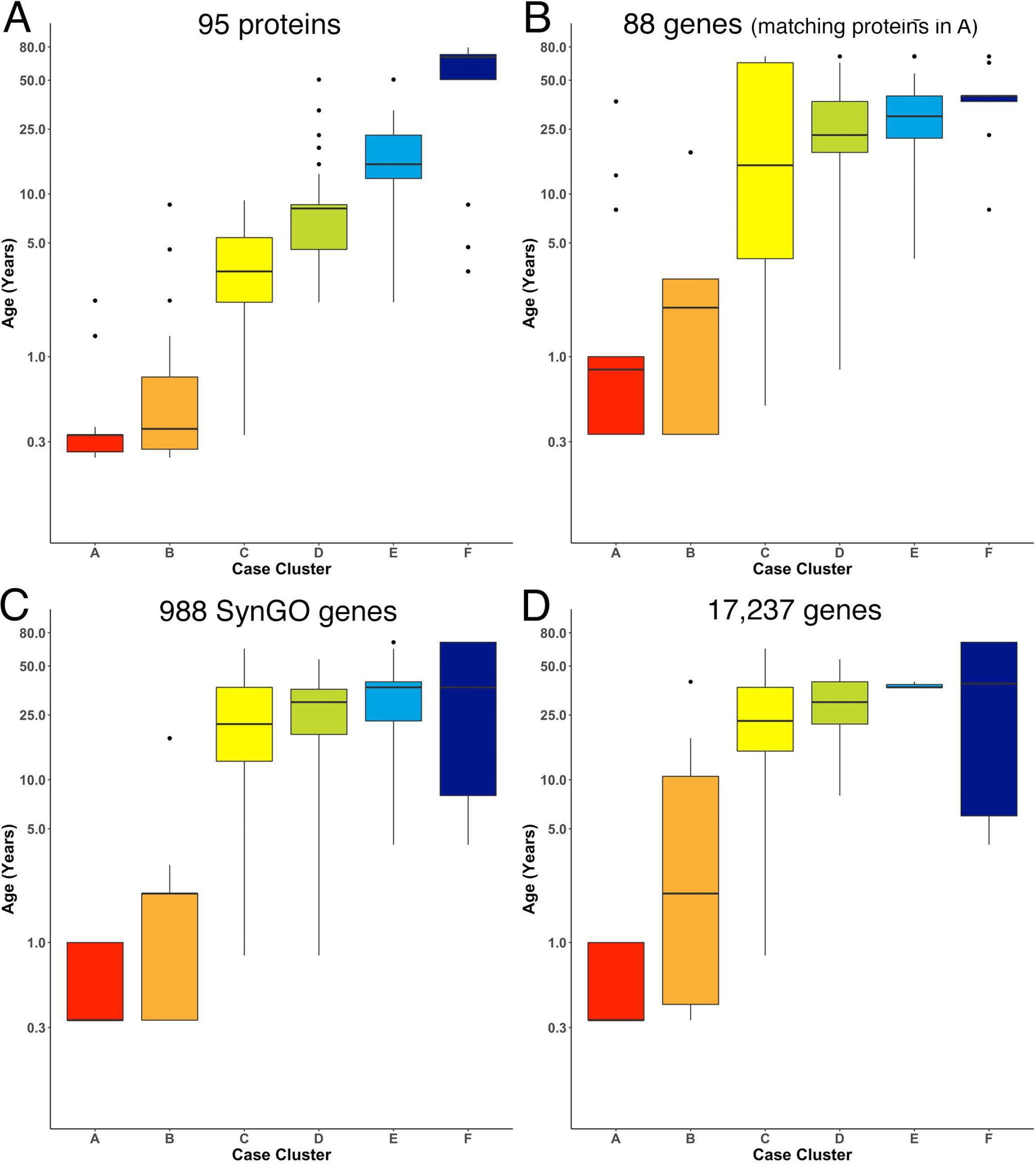
Age-related clusters for large numbers of proteins and genes. Expression data from A) 95 synaptic and immune-related proteins, B) 88 genes that correspond with the protein in panel A, C) 988 synaptic genes that correspond with the SynGO gene list and, D) 17,237 protein-coding genes was used to identify six age-related clusters. The cluster designation of each sample over 100 iterations of RSKC was used to visualize the distribution of sample ages. Boxplots denote the median and interquartile range of ages in each cluster, and points denote outliers.

Next, RSKC clustering was extended to the transcriptomic dataset from Kang et al. (2011) (Supplementary Material Table S2). First, RSKC was run using the 88 genes that matched the proteins in Fig. 7A. Even though the 2 datasets used different samples it was possible to compare the ages of the clusters because the range of ages and number of samples were similar. The progression of age-related clusters for the gene data (Fig. 7B) was similar to the protein clusters and there was a strong correlation (r=0.81) between the median ages of the 6 cluster pairs.

The strong correlation between the protein- and gene-cluster ages was particularly interesting because previous studies have shown that the correlation between large sets of protein and gene expression values is notoriously low (e.g. r ∼0.2) (Gry et al., 2009). To assess if the datasets used here simply had an unusually strong similarity between the lifespan changes in the expression values for each protein and gene pair we calculated those correlations. To facilitate this analysis the protein and gene expression values were normalized by calculating z-scores and the normalized values were partitioned into 6 age-bins (<1yr, 1-5yrs, 5-12yrs, 12-20yrs, 20-55yrs, >55yrs)(Supplementary Fig. S2). The correlation coefficient was then calculated for the 88 protein-gene pairs using the mean gene and protein expression values from the 6 age bins (Supplementary Fig. S3A). The mean correlation between the 88 protein-gene pairs was r=0.15 and the median correlation was only slightly higher (median r=0.21, 95% CI 0.04-0.25) (Supplementary Fig. S3B). Thus, it is unlikely that the strong correlation found between the ages of the protein- and gene-clusters arose from a simple linear relationship between those 2 types of molecular measurements. Instead, the common cluster ages for these different omics datasets suggest similar high dimensional patterns that RSKC uses to partition the samples into the series of age-related clusters.

Finally, we examined how well RSKC performed on datasets with measurements of hundreds to thousands of genes using 988 genes that overlap with the SynGO database of synaptic genes (Koopmans et al., 2019) and then with all 17,237 genes in the Kang dataset. The SynGO genes were analyzed to assess if a large set of functionally genes might reveal a different pattern of clusters from the full set of genes. The analysis of synaptic genes showed an age-related progression of the median age of the clusters (Fig. 7C). Compared with the protein clusters (Fig. 7A), the median age of the SynGO clusters jumped between clusters B and C (Fig. 7C) and a very similar pattern of age-related clusters was found when all 17,237 genes in the Kang dataset were used (Fig. 7D). Thus, RSKC cluster analysis of 95 proteins revealed the tightest age-related clusters, but the gene data also resulted in the partitioning of samples into age-related clusters. This finding contrasts with hierarchical clustering used by Kang et al. (2011)(Supplementary Fig. 8 in Kang et al 2011) that did not partition postnatal samples into age-related clusters. Thus, the optimization of sparse *K*-means cluster analysis (RSKC) for small sample sizes provides another approach for analyzing the human brain’s molecular development that is sensitive to the subtle molecular changes that occur across the postnatal lifespan.

### A note about selecting the number of clusters

An essential step in *k*-means clustering is selecting *k*, which denotes the number of groups to classify observation into. The correct choice of k is often ambiguous, as there are many different approaches for making this decision. Intuitively, an optimal k lies in between maximum generalization of the data using a single cluster and maximum accuracy by assigning each observation to its own cluster. One of the most common heuristics for determining k is the elbow plot method, where the sum of squared distances of observations to the nearest cluster center is plotted for various values of k. As k increases, the sum of squared distances tends towards zero. The “elbow” occurs at the point of diminishing returns for minimizing the sum of squared distances, and the k value at this point is selected as the optimal number of clusters (Thorndike. 1953).

To tailor the selection of k to RSKC, we applied the elbow method to the Weighted Within Sum of Squares (WWSS), the objective function maximized by the algorithm. WWSS was calculated for various values of *k* and averaged over 100 iterations. The elbow can be identified using the elbowPoint function in the *akmedoids* package (version 0.1.5) (Adepeju et al., 2020), which uses a Savitzky-Golay filter to smooth the curve and identify the x where the curvature is maximized. This method found that *k* = 6 was the optimal number of clusters for all of the applications of RSKC used in this paper.

There are more than 30 methods to determine the optimal values for *k* and a large number of journal papers (e.g. (Tibshirani et al., 2001)) and web resources (e.g<u>. Cluster Validation Essentials)</u> that can be used to learn more. The R packages *NbClust* (Charrad et al., 2015) and optCluster (Sekula, 2020) are particularly helpful tools for choosing the number of clusters because they test various methods for selecting *k* (Charrad et al., 2014).

### Application of RSKC clusters to study human visual cortex development

Previous studies using the datasets analyzed here (Murphy et al., 2005; Pinto et al., 2010, 2015; Williams et al., 2010; Kang et al., 2011; Siu et al., 2015, 2017) have examined molecular development by assigning samples into age-bins that approximate the lifespan stages defined by developmentalists. In contrast, the previous section describes a data-driven approach to partitioning samples into age-related clusters using sparse K-means clustering (RSKC). This use of unsupervised clustering raises the possibility that it might reveal aspects of human visual cortex molecular development that have escaped previous analyses. This section explores some of the information about human visual cortex development that can be revealed by examining the content of age-related clusters.

First, we compare partitioning of the samples into pre-defined age-bins versus data-driven clustering of the 23 proteins for post-mortem intervals (PMI), the proportion of cases, and the biological sex of the cases (Supplementary Fig. S4). The distribution of PMIs was similar between the two methods of partitioning the lifespan as was the proportion of samples and the balance of females and males in the bins. The progression of cluster ages was apparent when the age bins were colour-coded to reflect the cluster identity (Supplementary Fig. S4G). That histogram illustrated an interesting aspect of cortical development during young childhood (1-4 years) where samples in that age-bin were partitioned into 5 different clusters. Similar to previous studies that observed heightened childhood heterogeneity with waves of inter-individual variability that peak between 1-3 years (Pinto et al., 2015; Siu et al., 2017). The findings here suggest that the relationship between chronological and brain age varies across the lifespan.

The developmental trajectories of the 23 proteins were plotted using LOESS fits (95% CI) to the expression values (normalized to control), and each sample was colour-coded by their cluster assignment. The LOESS curves were ordered based on similar trajectories to illustrate the range of developmental patterns with some increasing (e.g. GABA_A_α1) or decreasing (e.g. GABA_A_α2) monotonically across the lifespan while others followed an inverted-U (e.g. gephyrin), an undulating pattern (e.g. VGAT) or remained relatively unchanged (e.g. GABA_A_α3)(Fig. 8A). The range of trajectories highlights the need for high-dimensional analyses to capture the complexity of this development. To help describe when the expression level of a protein in a cluster was above or below the overall mean, we implemented the over-representation analysis (ORA_phenotype function) described previously (Balsor et al., 2020)(Fig. 8B). Briefly, for each protein, a normal distribution was simulated using the mean and standard deviation of the expression values for all samples. Then the boxplots were colour-coded by comparing the expression values for each cluster with the simulated distribution. Here, the box for a cluster was coded as over-represented (red) if the 25th percentile was above 95% of the simulated distribution and under-represented if the 75th percentile was below 5% of the simulated distribution. Of course, other cutoff values for the ORA can be implemented to be more stringent or lenient for the colour-coding (e.g. Supplementary Fig. S5), or other methods such as estimation statistics (Bernard, 2019) can be used for this step depending on the nature of the question.

**Figure 8.**
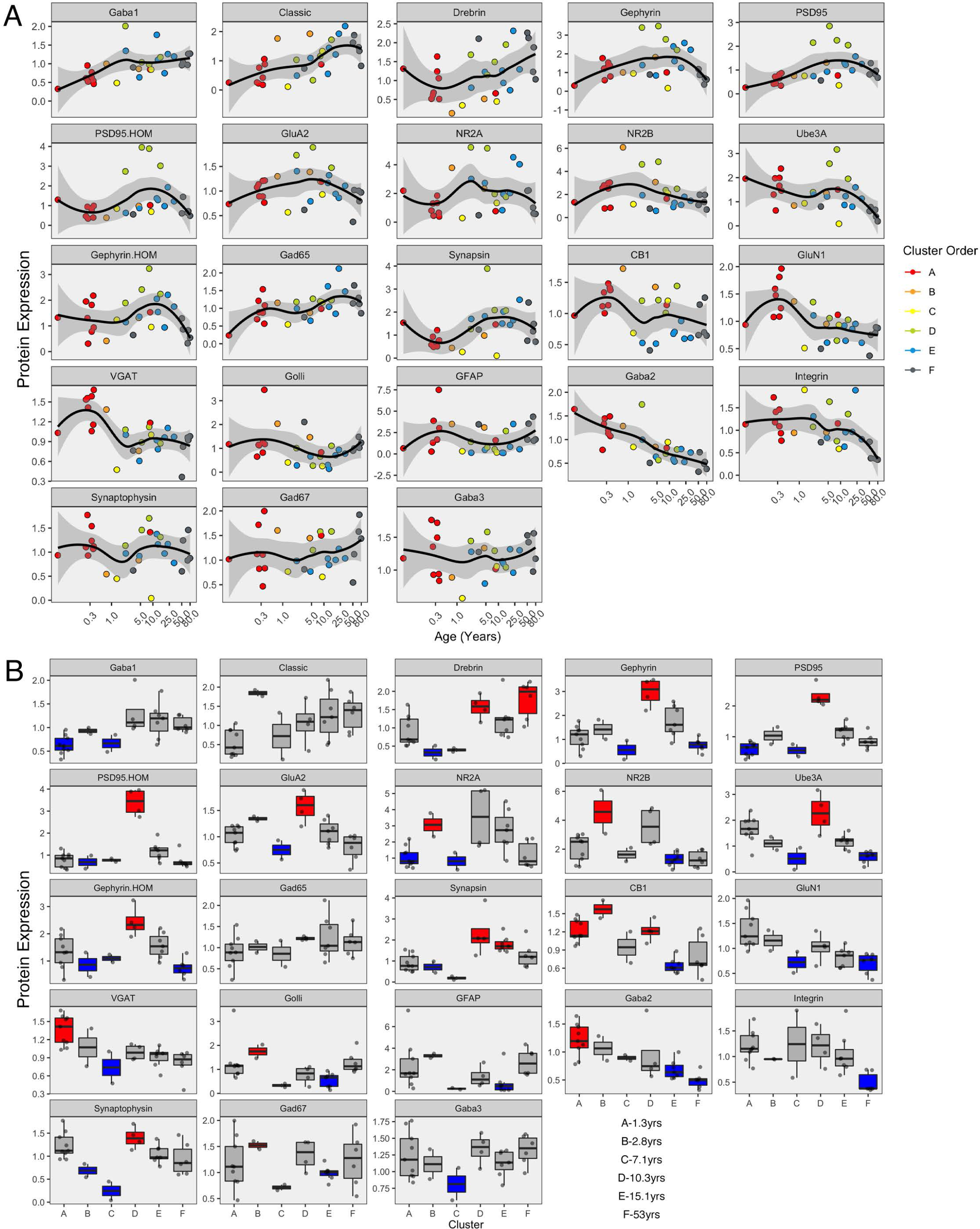
Development of proteins by age and by cluster. Expression profiles for each of the 23 proteins (A). Individual samples are represented by a single dot, and colored according to the corresponding RSKC cluster assignment. LOESS curves for each profile are shown in black, with 95% upper and lower confidence intervals shown bounding grey outline. Protein profiles are ordered according to similar developmental trajectories. **B)** Overrepresentation analysis showing protein expression as boxplots representing each cluster. Over-represented clusters were coloured red if the 25th percentile of the RSKC cluster was greater than the 95th percentile of a simulated normal distribution. Under-represented clusters were coloured blue if the 75th percentile of the RSKC cluster was less than the 5th percentile of a simulated normal distribution. Boxes that fell within the middle 90% of the simulated normal distribution were left grey.

Here, the ORA identified a range of over- or under-represented proteins in each cluster from a high of 12 proteins in clusters C to 5 proteins in cluster F (A - 6 proteins, B - 11 proteins, C - 12 proteins, D - 8 proteins, E - 7 protein, F - 5 proteins). These LOESS curves and boxplots for the expression of each protein help to describe development, but it is challenging to synthesize an overall pattern for human V1 development when confronted with making hundreds of pairwise comparisons. To address that problem we implemented a series of visualizations and analyses aimed at representing the high-dimensional nature of these data.

The first step in addressing the high-dimensional patterns of protein expression captured by the age-related clusters was to plot a bubble chart illustrating the expression levels of all 23 proteins for the 6 clusters. That visualization ordered the proteins by their RSKC weight and colour-coded each bubble with the normalized mean protein expression with blue representing low and red high expression levels (Fig. 9). The visualization helped identify that cluster D has high expression levels for many proteins. That cluster represents older children and the transition to adolescence (mean cluster age=10.3yrs, CI 9.6-11.1yrs) when rapid changes in cortical microstructure have been found (Norbom et al., 2021). In addition, groups of proteins with either high or low expression can be identified in a cluster, such as the higher expression of Golli-MBP, GFAP, CB1 and NR2B in cluster B. Thus, this visualization shows the mean expression for the 23 proteins in the 6 clusters, but it is still challenging to derive what differentiates the clusters. To address this, we applied our recently developed workflow (Balsor et al., 2019, 2020) that includes dimension reduction, identification of features and the construct of a plasticity phenotype visualization to characterize the development of the human visual cortex. This workflow is described in detail in a previous publication (Balsor et al., 2020).

**Figure 9.**
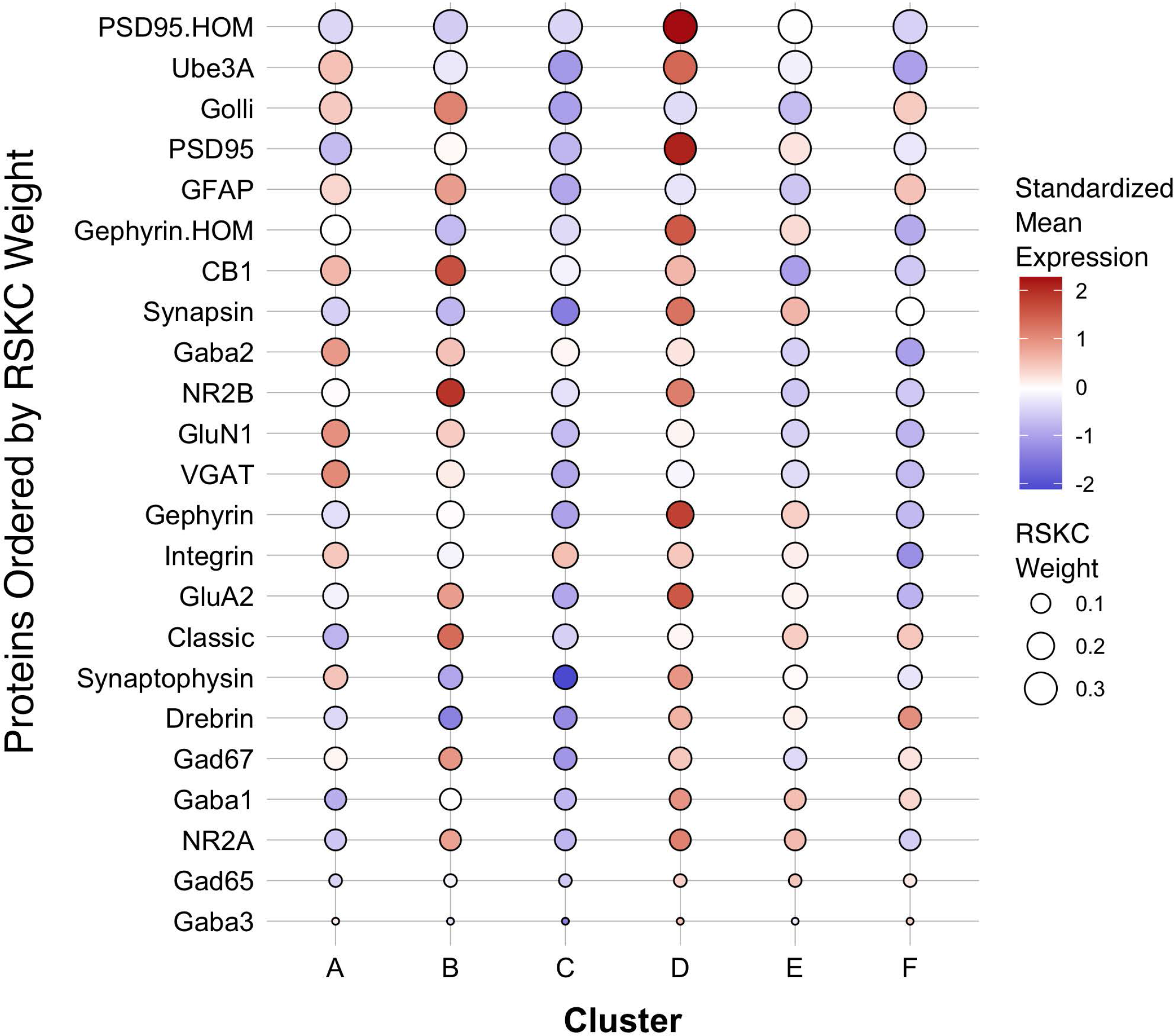
Bubble plot of mean protein expression across each cluster. Proteins are ordered by their corresponding RSKC weight with the highest weighted protein arranged at the top, and the lowest weighted protein at the bottom and ordered by developmental cluster from left to right. The color of the dot represents standardized protein expression for each cluster, while the size of the dot represents the RSKC weight (see legend)..

### Adding PCA for dimension reduction

This part of the workflow aims to reduce the dimensionality of the data by identifying combinations of functionally related proteins that we call features and using those features to capture the high dimensional pattern of brain development. The first step involves using Principal Component Analysis (PCA), a standard approach for reducing dimensionality when studying brain development (Jones et al., 2007; Beston et al., 2010; Bray, 2017). The scree plot showed that the first 3 dimensions capture ∼60% of the variance in the data (Supplementary Fig. S6), and the correlation matrix identified the strength of the relationship between each protein and the 23 dimensions. For example, the expression of Gephyrin and PSD95 was strongly correlated with Dim 1 while VGAT, GABA_A_α2, CB1 and GluN1 were strongly correlated with Dim 2. In addition, the quality of the representation for each protein on the first 3 dimensions was assessed using the cos^2^ metric. The cos^2^ (cosine square, coordinates square) conveys the quality of the representation of that variable using the projection angle onto each PC dimension. The closer that cos^2^ is to 1, the better the quality of that variable’s projection onto the dimension. The biplots illustrate the quality of the representation of each protein on Dim 1, 2 and 3 (Fig. 10 A&B) and show that some aspects of the RSKC-defined clusters are apparent when the samples are plotted in the PC space (Fig. 10 C&D). However, clustering of the samples in PC space was less distinct than illustrated in Figure 5 where tSNE plots were used to visualize clusters in the RSKC-weight transformed data.

**Figure 10.**
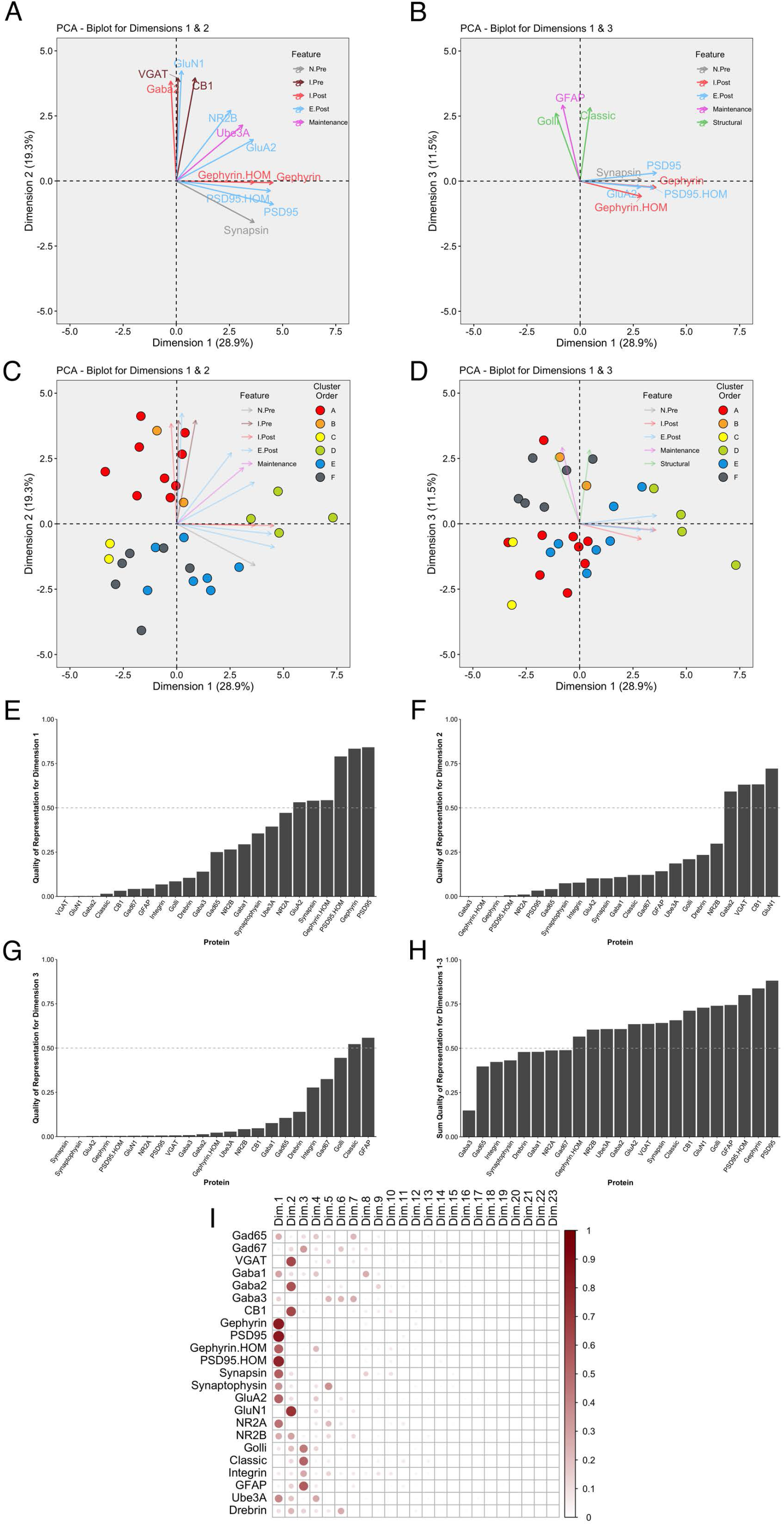
Examination of relevant PCA identified dimensions. PCA biplots **(A-D)** show protein features as vectors (arrows) and individual samples as dots on pairings of PCAdimensions 1 and 2 **(A,C)** and dimensions 1 and 3 **(B,D)**. The strength of the representation (cos^2^) for a protein on the given set of dimensions is reflected by the length of the vector, and only proteins with cos^2^ > 0.5 are shown. The colour of each point corresponds to their cluster, matching the original cluster colours in Figure 6. Bar plots represent the quality of representation of each protein with each dimension **(E-G)**, as well as the summed quality of representation across all three dimensions **(H)**. The dashed line represents cos^2^ cutoff value for representation of 0.5. **I)** Matrix illustrating the quality of representation for each protein with each PCA dimension, representing the strength (circle size) and direction (zero = white, positive = red) of cos^2^.

We examined which proteins were well represented by the first 3 dimensions by plotting the cos^2^ values for Dim 1, 2 and 3 (Fig. 10 E-G) and the sum of the cos^2^ for those dimensions (Fig. 10H). The matrix of cos^2^ values illustrated that only 2 of the proteins (GABA_A_α3 and Drebrin) were weakly represented by the first 3 dimensions. The remaining steps focus on Dim 1, 2 and 3 because they captured a large amount of the variance and had high-quality representations for most proteins.

### Comparing PCA and RSKC

We compared RSKC and PCA by assessing the similarity of RSKC weights and PCA cos ^2^ values for each protein (sum of Dim 1, 2 and 3) for the 23 proteins (Fig. 11A). There was a strong correlation (ρ = 0.72) between the 2 approaches; however, some proteins fell away from the line of best fit. Next, the differences between RSKC weights and PCA cos^2^ of the proteins were assessed using a Bland-Altman analysis (Giavarina, 2015). This used the calculated differences between the normalized measures and plotted those as the difference score for each protein. Also, interval bands were plotted to represent no difference between the RSKC and PCA measurements (blue band), when RSKC measurements were greater (positive red band) and when PCA measurements were greater (negative red band) (Fig. 11B). The blue band was slightly offset from zero, indicating a bias for the normalized PCA cos^2^ values to be greater than the RSKC weights. The plot identified key proteins, such as the Gephyrin and PSD95 homogenates, and Ube3A which were more strongly represented by the RSKC weights. All 3 of those proteins are essential molecular components that regulate the experience-dependent development of the visual cortex. For example, Ube3A is involved in the experience-dependent cycling of AMPA receptors (Greer et al., 2010), is required for ocular dominance plasticity (Yashiro et al., 2009; Sato and Stryker, 2010) and is selectively lost during ageing of the human visual cortex (Williams et al., 2010).

**Figure 11.**
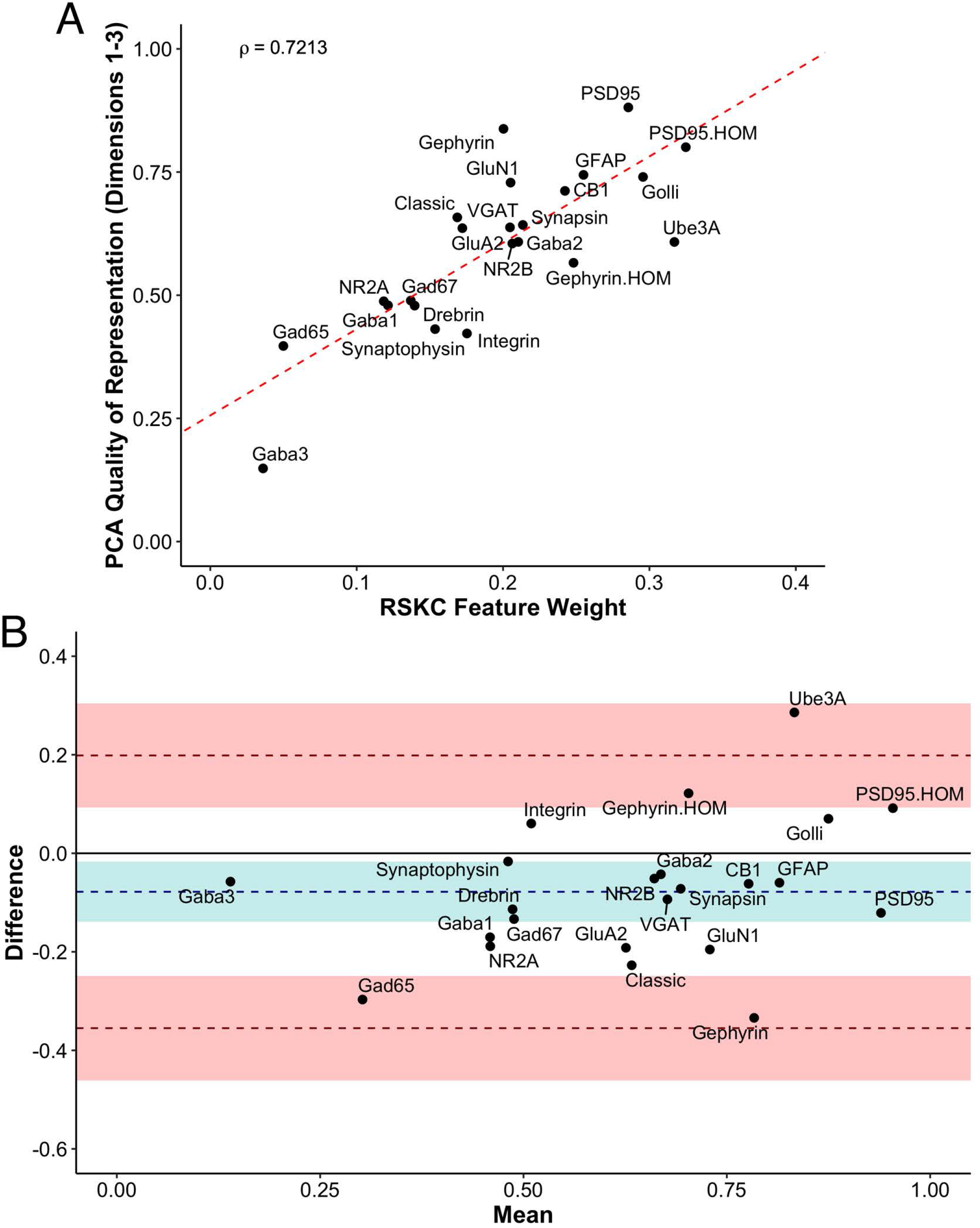
Exploring the relationship between PCA and RSKC feature identification. **A)** Scatter plot showing the PCA quality of representation (cos^2^) for the first 3 dimensions and the RSKC weights. The dashed line represents the line of best fit, and rho is Spearman’s rank correlation coefficient. **B)** Bland-Altman plot comparing PCA cos^2^ for the first 3 dimensions and RSKC weights for 23 proteins. The cos^2^ and RSKC weights were each computed as proportions of their respective maximum values. The dashed blue line represents the mean difference, with the 95% confidence intervals shown as the blue shaded area. The top dashed red line represents the upper limit of agreement (+1.96 SD) and the bottom dashed red line is the lower limit of agreement (–1.96 SD), with corresponding 95% confidence intervals shown as red shaded areas.

### Using PCA basis vectors to identify candidate plasticity features

The proteins in the dataset are known to regulate experience-dependent plasticity in the visual cortex (e.g. Quinlan et al., 1999a, 1999b; Fagiolini et al., 2003, 2004; Hensch, 2004, 2005; Hensch and Fagiolini, 2005; McGee et al., 2005; Philpot et al., 2007; Yashiro and Philpot, 2008; Cho et al., 2009; Gainey et al., 2009; Smith et al., 2009; Kubota and Kitajima, 2010; Larsen et al., 2010; Levelt and Hübener, 2012; Lambo and Turrigiano, 2013; Cooke and Bear, 2014; Guo et al., 2017; Turrigiano, 2017; Hensch and Quinlan, 2018). We took advantage of that *a priori* knowledge and the output from PCA to identify a new set of features that could be used to probe the neurobiology of the RSKC clusters. Although the RSKC weights reflect the contribution of individual proteins for partitioning the samples into clusters, the weights do not provide insights into combinations and balances of proteins that regulate plasticity. Thus, it is necessary to add another analysis that can help to identify those networks and balances of proteins that regulate experience-dependent plasticity.

This step is a semi-supervised approach to select combinations of proteins using the PCA output (cos^2^ values and the basis vectors) and the known functions of the proteins. These steps generate a list of candidate plasticity features that are combined to construct an extended phenotype (Dawkins, 1982). We call the collection of features a *plasticity phenotype* and it can be used to infer the plasticity state of the visual cortex. The approach is described in detail in Balsor et al (2020) and briefly outlined here.

Two heuristics were applied to identify combinations and balances among the proteins, using proteins that met the cos^2^ cutoff shown in Fig. 10H. First, the *a priori* knowledge about the function of the proteins in plasticity and development of the visual cortex was used to guide the inspection of the 3 basis vectors (Fig. 12A-C). Second, the amplitude and direction of each protein on the basis vector were used to select candidate features to sum or use in a relative difference index. For example, on PC1, we noted that 4 highly conserved synaptic markers (Pinto et al., 2015), the pre-synaptic proteins synapsin and synaptophysin and the post-synaptic proteins PSD95 and gephyrin had large positive amplitudes, so they were summed to create one of the candidate features (PGSS). On PC2, the receptor subunits GABA_A_α1 and GABA_A_α2 had opposite directions, so these were used for an index (GABA_A_α1:GABA_A_α2) because the balance between those subunits is developmentally regulated and governs the kinetics of the GABA_A_ receptor (Gingrich et al., 1995; Bosman et al., 2002; Heinen et al., 2004; Hashimoto et al., 2009). Finally, on PC3, we noted that GFAP and integrin had the largest amplitudes, and they were in opposite directions. Those 2 proteins are expressed by astrocytes, and the expression of integrin receptors is increased on reactive astrocytes (Lagos-Cabré et al., 2020), so an index was calculated (GFAP:Integrin). Applying the heuristics resulted in 13 candidate features, including 5 protein sums identified using the basis vector for PC1 and 8 indices from PC2 and PC3 (Fig. 12D). The features were validated by calculating each feature using the expression values for the 23 proteins (Supplementary Material Table 6) and correlating those with the eigenvalues for the 3 PC dimensions (Fig. 12D).

**Figure 12.**
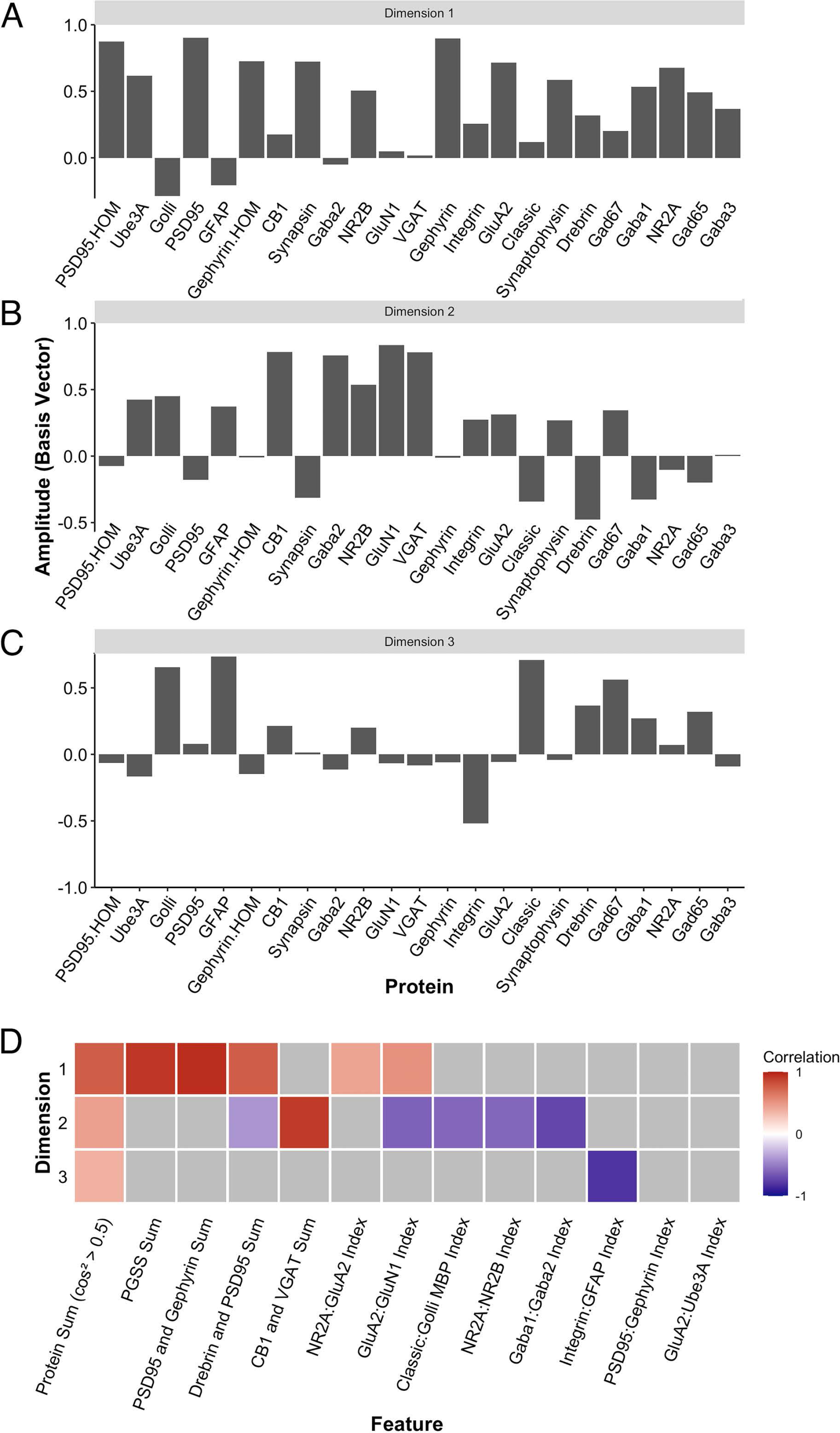
Candidate feature identification using principal component analysis. Histograms showing the amplitude of the basis vector for each protein across **A)** dimension 1, **B)** dimension 2, **C)** dimension 3. **D)** The correlations between the protein sums or indices and the first 3 PCA dimensions. Non-grey cells represent significant correlations after Bonferroni correction, with the colour indicating the magnitude and direction of the correlation (negative = blue, zero = white, positive = red).

LOESS curves and boxplots were made for all of the candidate features to illustrate how they changed across the lifespan and identify if features were over- or under-represented in a cluster (Fig. 13 A&B, Supplementary Fig. S7). One aspect of development apparent in the boxplots was the over-representation of the protein sums in cluster D. That cluster has a mean age of 10.3 years (SD 8.4 years), which corresponds with the end of the critical period for developing amblyopia in children (Lewis and Maurer, 2005; Birch, 2013) and a stage of human cortex development often described by synaptic exuberance, growth and changing state of plasticity. Furthermore, animal research has shown that excess excitation (Fagiolini and Hensch, 2000; Fagiolini et al., 2004) and expression of proteins regulating that activity, especially PSD95 (Huang et al., 2015), can close the critical period.

**Figure 13.**
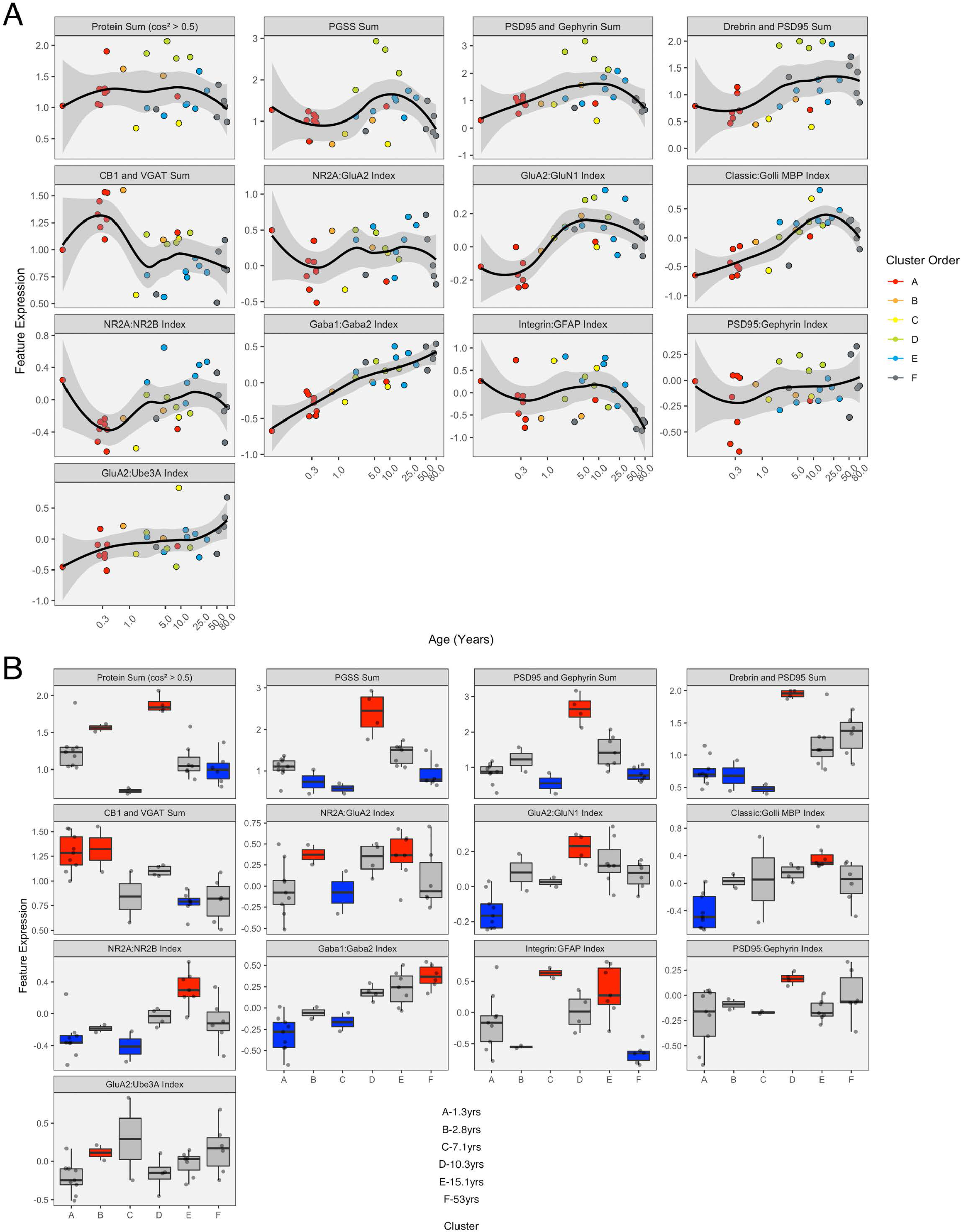
Expression of extracted features with respect to age and by cluster. **A)** LOESS trajectories illustrating the expression of protein sums and indices. Points are coloured corresponding to the clusters in Figure 8, and the 95% confidence intervals around each curve are coloured in grey. **B)** Boxplots show the expression of each feature for the 6 clusters. A simulated normal distribution was sampled to obtain 5th and 95th percentile values. Boxes were coloured red (i.e. over-represented) if the 25th percentile of the feature cluster was greater than the 95th percentile of the normal distribution. Boxes were coloured blue (i.e. under-represented) if the 75th percentile of the feature cluster was less than the 5th percentile of the simulated distribution. Otherwise, boxes were coloured grey.

### Analyzing plasticity phenotypes for the RSKC clusters

Finally, the 11 features with significant correlations were used to construct a plasticity phenotype that was combined with the 6 clusters. A correlation matrix was made using the values for the features calculated from the protein expression for each sample (Table 2). The matrix and surrounding dendrogram showed that the protein sum and indices were separated into different tree branches. The order of the features in the correlation matrix was used for the bands in the plasticity phenotype visualization. In the phenotype, the median of each feature was represented as a colour-coded band for the 6 clusters (Fig. 14B). Together, the 66 colour-coded feature bands captured the high dimensional pattern of neurobiological changes across the lifespan. The protein sums represented by grey levels convey a pattern with specific groups of proteins that are highly expressed early in development (clusters A & B) and a broad wave of expression in older childhood (cluster D). The indices reflect the multiple timescales of molecular development that are the hallmarks of the human visual cortex (Siu and Murphy, 2018). However, even with undulating features and different timescales, all appear to arrive at a similar level of maturation in cluster E.

**Figure 14.**
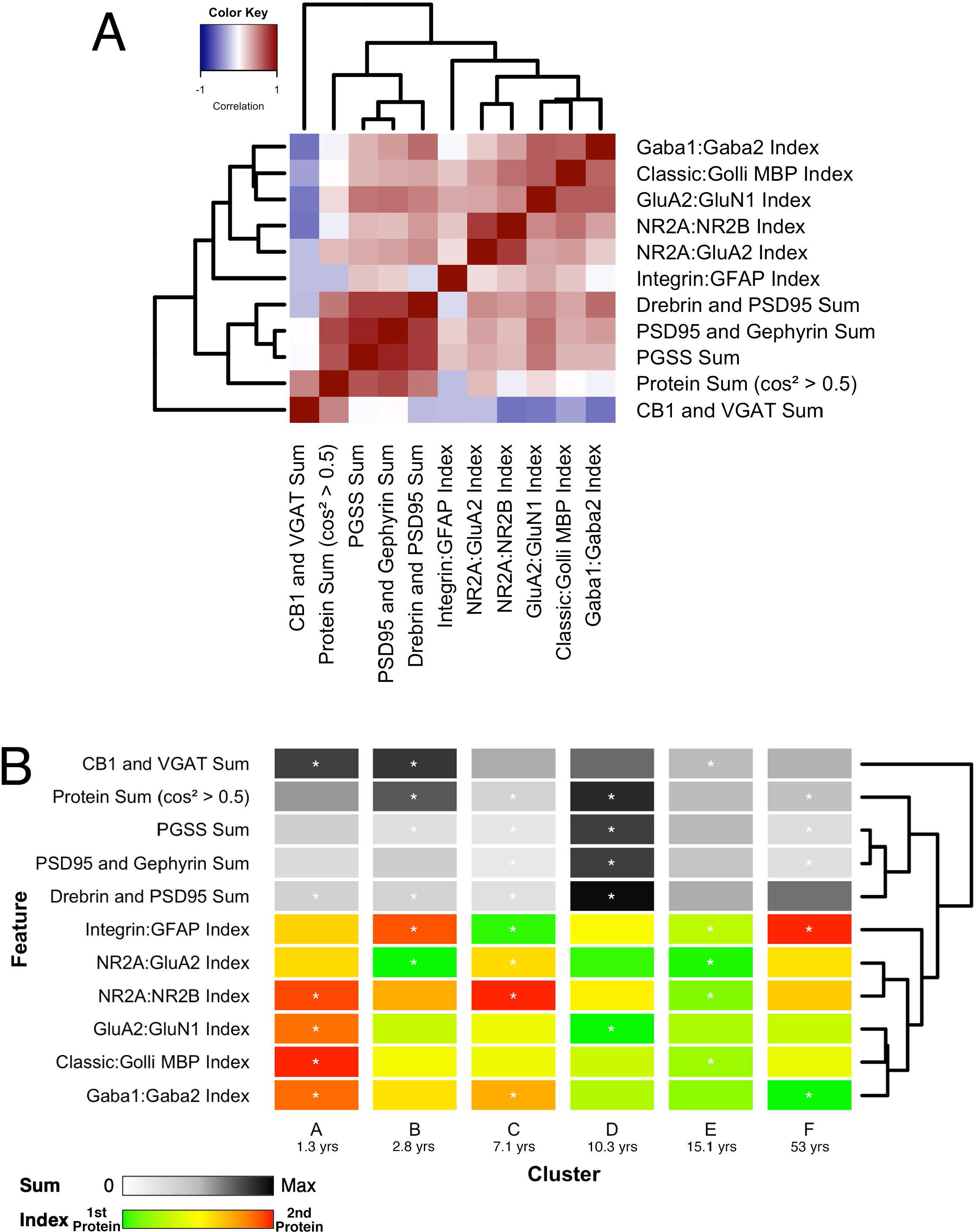
Associations between selected features and feature phenotype by cluster. **A)** Correlation heat map between protein sums and indices, with strength and direction of Pearson’s R correlation represented by the colour (negative = blue, zero = white, positive = red), and arranged by similar pairwise correlations using a wrapped dendrogram. Features were selected if they were significantly correlated with any of the first three PCA dimensions. **B)** The plasticity phenotype was visualized using colour-coded horizontal bars representing the median expression of selected features across clusters. For protein sums, the colour ranges from white (zero) to grey (midpoint) to black (maximum protein sum across all features). For the protein indices, the colour ranges from green (favouring the first protein in the index) to yellow (balance of the 2 proteins) to red (favouring the second protein in the index). Asterisks indicate features that were found to be either over- or under-represented. The features are arranged according to the same dendrogram generated in A.

Combining the features and clusters into a visualization simplified this complex dataset and facilitated linking the clusters with sets of neurobiologically meaningful features. The asterisks on the feature bands indicate the ones identified as over- or under-represented in Fig. 13B. Each cluster had a unique group of features that deviate from the average, and those represent the neurobiological mechanisms that differentiate the age-related clusters. For example, the set of 4 red bands for the young visual cortex (cluster A) was unique and showed that the indices were dominated by the NMDA receptor subunits NR2B and GluN1, the Golli family of myelin basic protein (MBP) and the GABA_A_α2 receptor subunit. In contrast, the older visual cortex (cluster F) was distinguished by a set of 3 light grey protein sum bands, a red band indicating more GFAP and a green band indicating more GABA_A_α1. Finally, the overall appearance of the protein sums and indices for cluster D gives the impression of a transition stage in the development of the visual cortex when exuberant protein expression (dark grey bands)(Huang et al., 2015) and the shift in protein balances (green bands)(Quinlan et al., 1999a, 1999b; Fagiolini and Hensch, 2000; Chen et al., 2001; Philpot et al., 2001; Fagiolini et al., 2003, 2004; Hensch, 2005; Hall and Ghosh, 2008; Smith et al., 2009) signals the end of the critical period.

A raincloud plot of the samples in the 6 clusters shows the range of ages that correspond with the plasticity phenotypes (Fig. 15). The distribution of sample ages in the clusters appears like a series of overlapping waves extending beyond the ages of the traditional pre-defined age-bins included in Fig. 15 as vertical lines. For example, for cluster D, the wave’s peak falls into the age-bin associated with the end of the period for developing amblyopia (5-12 years). However, cluster D also includes younger and older samples suggesting that the end of the sensitive period may not occur uniformly among individuals. Furthermore, other clusters overlap the 5-12-year-old age-bin suggesting that multiple phenotypes can be found during certain age-bins. Thus, the cluster analysis helped reveal aspects of visual cortex development that are obscured by using pre-defined age bins, which is that chronological- and brain-age often diverge (Cole et al., 2019).

**Figure 15.**
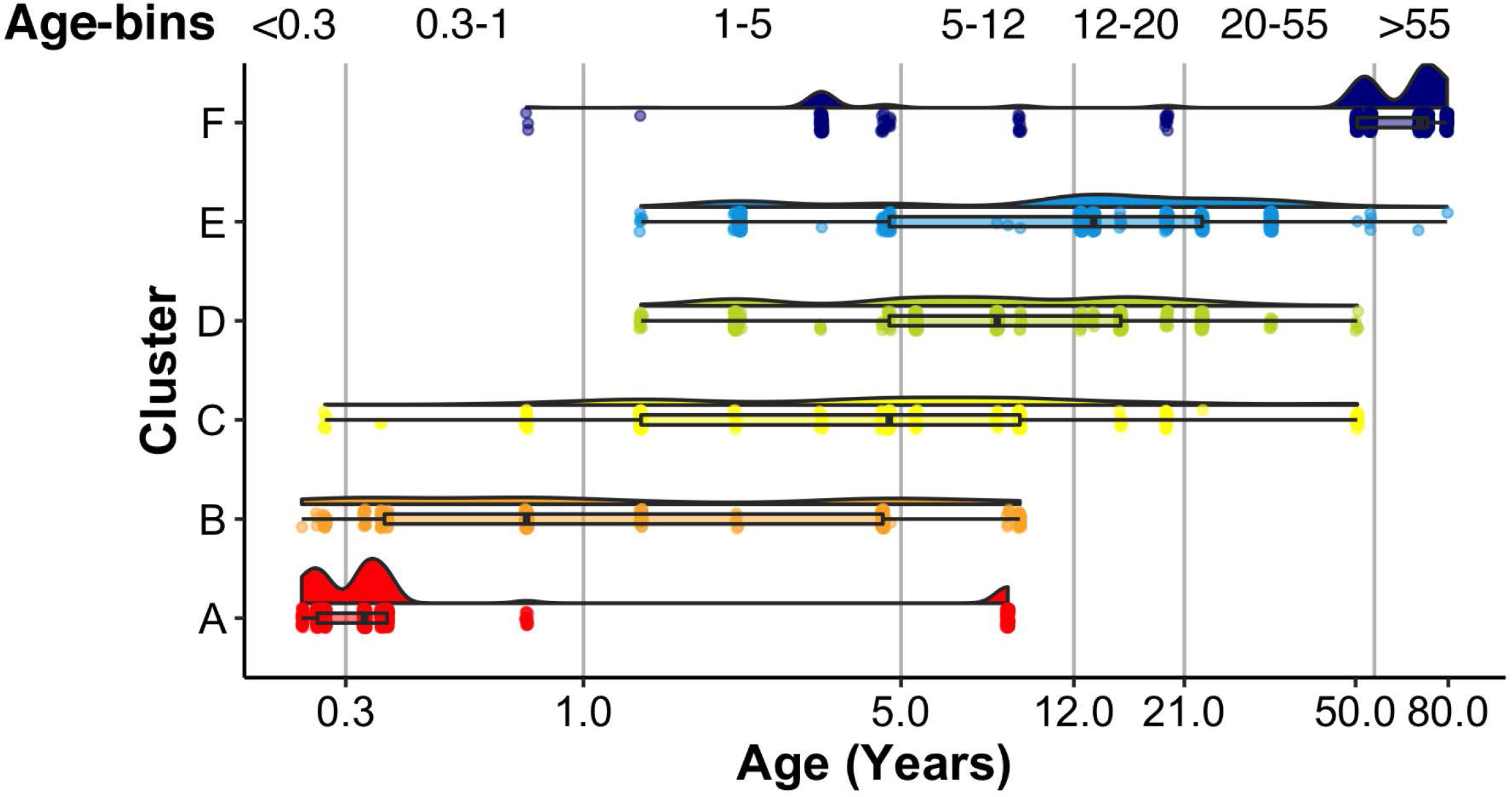
Raincloud plot showing the distribution of sample ages by cluster. Vertical lines correspond to the bounds of pre-defined age bins. Samples corresponding to the youngest age (i.e. 20 days) are not shown.

## Discussion

The current study shows that the application of sparse clustering leverages the high dimensional nature of proteomic and transcriptomic data from human brain development to find age-related clusters that are spread across the lifespan. In particular, robust sparse *K*-mean clustering (RSKC) using measurements of proteins or genes from the human visual cortex partitioned samples into clusters that progressed from neonates to older adults. The iterative reweighting of the measurements to focus on the proteins or genes that carry the most information about lifespan changes led to robust age-related clustering of the data. Furthermore, especially for the datasets focusing on 95 proteins or genes, the clusters represented early development, young childhood, older childhood, adolescence and adulthood. Thus, sparse clustering provides a robust approach for identifying proteomic or transcriptomic defined brain ages that overlap with behavioural and brain imaging findings of gradual and prolonged human brain maturation.

Many factors come into play when selecting an appropriate clustering algorithm for a study. Here, we considered the goal of the study (to resolve sometimes subtle age-related changes in molecular mechanism), the structure of the dataset ( p∼n to p>>n), and the output of the algorithm (is it just the clusters or is feature selection included). Sparse K-means clustering was selected because it fit all of those considerations. We know from previous studies of the molecular development of the human brain that there can be subtle differences between age groups (refs), and yet even small changes in protein or gene expression will alter neural function. Therefore, we looked for algorithms designed for omics datasets where subtle changes in a subset of the genes or proteins would identify important characteristics of the data. The development of sparse K-means clustering by Witten and Tibshirani (ref) was partially inspired by the need to better cluster a breast cancer dataset. In that dataset, subtle differences in gene expression significantly impacted patient outcomes, but standard clustering approaches did not pick those up. In addition, sparse clustering was developed to address datasets, like ours and the breast cancer data where the structure is *p∼n* to *p>>n*. Sparse K-means clustering is a good fit for those high dimensional structures because it minimizes the within-cluster sum of squares with a dissimilarity measure while maximizing the between-cluster sum of squares by iteratively reweighting the measures. Finally, and most importantly, sparse K-means clustering performs feature selection. The examples in this paper show the reweighted proteins and those distributions identifying how much each protein contributes to partitioning the samples into clusters. That matrix is sparse, with unimportant proteins having near-zero weights and important ones having non-zero weights. Those weights are essential for cluster analysis to help with making neurobiologically relevant interpretations of brain development from the cluster analysis.

Various other algorithms, including linear and non-linear dimension reduction (e.g. tSNE, MDS, PCA), can separate developmental samples. In this paper, we found that both tSNE and PCA show some age-related progress in the arrangement of the samples. Also, the original Kang paper (Kang et al., 2011) used multidimensional scaling (MDS) to separate the samples across MDS 1 and 2. Then the points were colour-coded by pre-defined age bins to show a left to right flow from early prenatal to older adults. However, it was not apparent which genes mapped on to those dimensions. The selection of features in the form of the weights is a key difference between sparse K-means clustering and standard clustering approach that was critical for the current study.

The current study is not exhaustive of clustering approaches, as the number of unsupervised clustering algorithms for analyzing high dimensional data is rapidly expanding. For example, new sparse clustering algorithms include innovation at the level of the lasso-type penalty used to adjust observation weights (Brodinová et al., 2019). Accordingly, the “best” algorithm for understanding molecular brain development will continue to change as new approaches are developed. Rather than acting as a prescriptive guide for which algorithm to use, the current study highlights the challenges raised when applying high dimensional clustering to studies using postmortem brain samples. In particular, developmental studies that use postmortem human brain tissue often have more measurements than samples (*p* > *n*) and require clustering algorithms optimized for high dimensional data structures. The examples showed that the RSKC algorithm worked well for a wide range of observations (*p*) from 7 to 17,237. However, the age-related progression of the 95 proteins and 88 gene datasets (Fig. 7A&B) were more distinct than the clustering using 988 SynGO or the full 17,237 gene dataset (Fig. 7 C&D).

The succession of age-related clusters found for the visual cortex aligns with some critical milestones in visual development. Using measurements of molecular mechanisms that regulate experience-dependent plasticity, the clusters illustrated in Figure 5 show that Cluster A overlaps the start of the sensitive period for binocular vision at 4-6 months and Cluster B the peak of that sensitive period at 1-3 years (Banks et al., 1975). Furthermore, Cluster D aligns with the maturation of contrast sensitivity (Ellemberg et al., 1999), motion perception (Ellemberg et al., 2002), and the end of the period for the susceptibility of developing amblyopia (6 -12 years) (Epelbaum et al., 1993; Keech and Kutschke, 1995; Lewis and Maurer, 2005). The oldest cluster, Cluster F, highlights ages when cortical changes reduce performance on several visual tasks (Owsley, 2011). The alignment with visual milestones suggests that the clusters might provide insights into the molecular mechanisms that regulate various aspects of visual development and visual function dynamics across the lifespan. Notably, the molecular mechanisms are well studied in animal models. Thus, this information for the human cortex may be seen as a bridge linking results from animal studies with human neurobiology that can help interpret brain imaging and visual perception findings.

By combining the RSKC clustering with PCA, we identified plasticity-related features and constructed a plasticity phenotype that was applied to each cluster (Fig. 14). The term *plasticity phenotype* has been used before to describe the waxing and waning of gene expression in the developing brain (Smith et al., 2019). Here we used the term to describe an extended phenotype (Dawkins, 1982) because the proteins in the dataset have known functions in regulating experience-dependent plasticity in the visual cortex (Quinlan et al., 1999a, 1999b; Fagiolini et al., 2003, 2004; Hensch, 2004; Hensch and Fagiolini, 2005; McGee et al., 2005; Philpot et al., 2007; Yashiro and Philpot, 2008; Cho et al., 2009; Gainey et al., 2009; Smith et al., 2009; Kubota and Kitajima, 2010; Larsen et al., 2010; Levelt and Hübener, 2012; Lambo and Turrigiano, 2013; Cooke and Bear, 2014; Guo et al., 2017; Turrigiano, 2017; Hensch and Quinlan, 2018). Thus, the plasticity phenotype can be used to infer the potential for experience-dependent plasticity in the different clusters and provide a new perspective on the maturation of the human visual cortex.

Each cluster had a unique set of features that were over- or under-represented in the plasticity phenotype, and those features were apparent in the phenotype visualization. Notably, the features were selected using a semi-supervised approach with a series of heuristics that included protein combinations and balances known to regulate experience-dependent plasticity. As a result, the unique sets of features can be compared with the literature to infer the likely state of experience-dependent plasticity for a cluster. For example, balances in the youngest cluster (A) were dominated by receptors that are known to facilitate experience-dependent plasticity in the visual cortex (Kleinschmidt et al., 1987; Quinlan et al., 1999a, 1999b; Philpot et al., 2001; Fagiolini et al., 2003, 2004; Iwai et al., 2003; Hensch, 2004; Cho et al., 2009; Jiang et al., 2010). In contrast, cluster D overlaps the end of the period of susceptibility to develop amblyopia, and has peaks in protein expression, especially PSD95 that are known to close the critical period in animal models (Huang et al., 2015). The features in cluster D also appeared to mark the transition from juvenile features found in clusters A, B and C to the mature and ageing patterns in clusters E and F. Moreover, the range of ages in a cluster appeared as a series of overlapping waves in the raincloud plot, thereby illustrating that chronological- and brain-age have a complex relationship.

Clustering the data collected from human postmortem tissue samples to reveal the age-related progression in the brain’s molecular complexity is just the start of using high dimensional analyses. The application of modern exploratory data-driven approaches reveals novel aspects of human brain development, such as the risk for mental illness (Li et al., 2018) or divergence from other primates (Zhu et al., 2018). Identifying an appropriate high dimensional clustering technique opens the door to many other downstream analyses to interrogate different clusters’ molecular makeup. A critical benefit of clustering with RSKC is that it outputs the feature weights. Those weights reveal the impact of specific proteins or genes on differentiating the brain’s molecular environment during the progression of lifespan stages. Those proteins and genes can be used as the input to gene ontology (GO) analysis to catalogue the molecular processes, cellular components, and biological processes that dominate the stages. Or the opposite can be done as shown in the paper where the 988 genes corresponding to the SynGO database were used to cluster the samples. The clusters can also be used for differential gene expression analysis to highlight which features are enriched during various lifespan stages. For example, the top-weighted molecular features from the RSKC analysis may be useful for creating a phenotype that provides a biologically meaningful characterization of the high dimensional changes that occur in different stages of the lifespan (Balsor et al., 2020).

An interesting finding of the current study is the overlapping ages among the clusters. While this may be viewed as imperfect partitioning of samples by the clustering algorithms, it may also reflect the human brain development’s true heterogeneity. In other words, developmental periods may not necessarily be described by a single omic phenotype. Instead, the classically defined developmental stages may be characterized by two or more distinct patterns of gene or protein expression in the brain. This molecular heterogeneity may shed light on findings such as the substantial inter-individual variation in cortical responses measured by fMRI studies in infants (Born et al., 2000). Also, the overlapping ages among clusters may reflect periods of stationary fluctuations in the brain’s developmental trajectory, representing transitions from one molecular state to the next, similar to language development models (Sanchez-Alonso and Aslin, 2020).

Addressing how human brain development proceeds is an important question that will require large amounts of new data and algorithms that capture the local and global structure of high dimensional trajectories, including ones with gradual noisy changes and non-linear transitions. One approach could include repeated MRI measurements during the ages that overlap among molecular clusters to assess if those ages have heightened intra- or inter-individual variation in brain responses. Those studies will help identify ages during development with gradual but noisy change from ages with non-linear transitions in the gene and protein expression pattern in the developing human brain. Ultimately, the models will need to include multi-omics data and link with brain imaging to understand how the human brain develops fully.

## Conclusions

The last decade has seen remarkable growth in the number of studies examining the human brain’s molecular features. In parallel, high throughput tools have dramatically increased the amount of data collected for every sample. The complexity and high dimensional nature of those datasets have spurred the need for more guidance in selecting appropriate tools to analyze those big data. Some studies are now collecting data from 100s or 1000s of human brain postmortem samples (e.g. PsychENCODE), but studies of development still have many fewer tissue samples, and the ages of the cases are spread across the lifespan. The small sample sizes of the developmental datasets make it difficult to apply many commonly used high dimensional clustering methods. Those methods lack the sensitivity needed to reveal robust clusters defined by the subtle differences in genes or proteins that occur across the postnatal lifespan. At the same time, sparsity-based clustering algorithms designed for small sample size have emerged. In this guide, we explored the application of sparsity-based clustering and showed that one algorithm, Robust Sparse *K*-means Clustering, is a good fit for revealing the subtle and gradual changes of human brain development that occurs from birth to aging. In the next decade, the amount of data collected from each postmortem brain sample will only continue to grow as single-cell RNA sequencing methods are applied to studying human brain development. Furthermore, the push to integrate multimodal measurements, from molecules to imaging of human brain development will heighten the demand for robust high dimensional analysis tools. Neuroscientists will continue to face many challenges identifying rigorous methods to analyze those sparse and very high dimensional datasets. Nevertheless, careful selection of high dimensional analytical techniques that are designed for small sample sizes can be expected to have an impact on the discovery of novel aspects of human brain development.

## Authors’ contributions

JB Designed research, performed research, analyzed data, wrote/revised the paper; KA designed research, analyzed data, wrote/revised the paper; DS performed research, analyzed data, wrote/revised the paper; RK performed research, analyzed data, revised the paper; JZ performed research, analyzed data, revised the paper; EJ analyzed data, revised the paper; KM designed research, analyzed data, wrote/revised the paper.

## Acknowledgements

We thank Brendan Kumagai for help with coding. Previous studies from our lab collected the proteomic data, and the human postmortem tissue samples were obtained from the NIH NeuroBioBank (Pinto et al., 2010, 2015; Williams et al., 2010; Siu et al., 2015, 2017). The transcriptomic data were from (Kang et al., 2011) (<u>GSE25219</u>).

## Data and Code Availability

The data used to support the findings in this manuscript and code are available here: https://osf.io/6vgrf/

