## Supplementary Tables_Figures for "A Practical Guide to Sparse K-Means Clustering for Studying Molecular Development of the Human Brain"

**Supplementary Material – Tables and Figures**

Justin L. Balsor[1], Keon Arbabi[1], Desmond Singh[2], Rachel Kwan[2], Jonathan Zaslavsky[2], Ewalina Jeyanesan[1], Kathryn M. Murphy[1,2]

[1]McMaster Integrative Neuroscience Discovery and Study (MiNDS) Program, McMaster University, Hamilton, ON, L8S 4K1, Canada

[2]Department of Psychology, Neuroscience & Behavior, McMaster University, Hamilton, ON, L8S 4K1, Canada

**Supplementary Table 1**. *Human samples from Murphy lab studies*. The age, age group, biological sex, and post-mortem interval (PMI) for each of the human cortical tissue samples is listed.

| Age | Age Group | Sex | PMI (Hours) |
|---|---|---|---|
| 20 days | Neonate | M | 9 |
| 86 days | Neonate | F | 23 |
| 96 days | Neonate | M | 12 |
| 98 days | Neonate | M | 16 |
| 119 days | Neonate | M | 22 |
| 120 days | Neonate | M | 23 |
| 133 days | Infant | M | 16 |
| 136 days | Infant | F | 11 |
| 273 days | Infant | M | 10 |
| 1 year 123 days | Young Children | M | 21 |
| 2 years 57 days | Young Children | F | 21 |
| 2 years 75 days | Young Children | F | 11 |
| 3 years 123 days | Young Children | F | 11 |
| 4 years 203 days | Young Children | M | 15 |
| 4 years 258 days | Young Children | M | 17 |
| 5 years 144 days | Older Children | M | 17 |
| 8 years 50 days | Older Children | F | 20 |
| 8 years 214 days | Older Children | F | 20 |
| 9 years 46 days | Older Children | F | 20 |
| 12 years 164 days | Teens | M | 22 |
| 13 years 99 days | Teens | M | 5 |
| 15 years 81 days | Teens | M | 16 |
| 19 years 76 days | Teens | F | 16 |
| 22 years 359 days | Young Adults | M | 4 |
| 32 years 223 days | Young Adults | M | 13 |
| 50 years 156 days | Young Adults | M | 8 |
| 53 years 330 days | Young Adults | F | 5 |
| 69 years 110 days | Older Adults | M | 12 |
| 71 years 333 days | Older Adults | F | 9 |
| 79 years 181 days | Older Adults | F | 14 |

**Supplementary Table 2**. *Human samples from Kang et al. (2011)*. Samples for the primary visual cortex (V1C) are shown. Age is either postnatal months (PMonth) or years (Y).

| ID | Brain Code | Region | Sex | Age | Years | Weeks | PMI | Hemisphere |
|---|---|---|---|---|---|---|---|---|
| GSM704398 | HSB121 | V1C | M | 4 PMonth | 0.33 | 17 | 9.5 | L |
| GSM704399 | HSB121 | V1C | M | 4 PMonth | 0.33 | 17 | 9.5 | R |
| GSM704607 | HSB132 | V1C | M | 4 PMonth | 0.33 | 17 | 22 | L |
| GSM704608 | HSB132 | V1C | M | 4 PMonth | 0.33 | 17 | 22 | R |
| GSM704729 | HSB139 | V1C | M | 4 PMonth | 0.33 | 17 | 20 | L |
| GSM704577 | HSB131 | V1C | F | 6 PMonth | 0.5 | 26 | 26 | L |
| GSM705106 | HSB171 | V1C | M | 10 PMonth | 0.83 | 43 | 18 | L |
| GSM704430 | HSB122 | V1C | F | 1 Y | 1 | 52 | 18 | L |
| GSM704431 | HSB122 | V1C | F | 1 Y | 1 | 52 | 18 | R |
| GSM704802 | HSB143 | V1C | F | 2 Y | 2 | 104 | 12 | L |
| GSM704803 | HSB143 | V1C | F | 2 Y | 2 | 104 | 12 | R |
| GSM705120 | HSB172 | V1C | M | 3 Y | 3 | 156 | 16 | L |
| GSM705133 | HSB173 | V1C | F | 3 Y | 3 | 156 | 8 | L |
| GSM704353 | HSB118 | V1C | M | 4 Y | 4 | 208 | 20 | R |
| GSM704745 | HSB141 | V1C | M | 8 Y | 8 | 416 | 30 | L |
| GSM705145 | HSB174 | V1C | M | 8 Y | 8 | 416 | 16 | L |
| GSM705159 | HSB175 | V1C | F | 11 Y | 11 | 572 | 22 | L |
| GSM704492 | HSB124 | V1C | F | 13 Y | 13 | 676 | 19.5 | R |
| GSM704368 | HSB119 | V1C | M | 15 Y | 15 | 780 | 14.5 | L |
| GSM704151 | HSB105 | V1C | M | 18 Y | 18 | 936 | 28 | L |
| GSM704152 | HSB105 | V1C | M | 18 Y | 18 | 936 | 28 | R |
| GSM704545 | HSB127 | V1C | F | 19 Y | 19 | 988 | 9.5 | L |
| GSM704561 | HSB130 | V1C | F | 21 Y | 21 | 1092 | 18 | L |
| GSM704775 | HSB142 | V1C | M | 22 Y | 22 | 1144 | 18 | L |
| GSM704776 | HSB142 | V1C | M | 22 Y | 22 | 1144 | 18 | R |
| GSM704711 | HSB136 | V1C | M | 23 Y | 23 | 1196 | 10.5 | L |
| GSM704712 | HSB136 | V1C | M | 23 Y | 23 | 1196 | 10.5 | R |
| GSM704619 | HSB133 | V1C | F | 27 Y | 27 | 1404 | 21 | L |
| GSM704527 | HSB126 | V1C | F | 30 Y | 30 | 1560 | 9.5 | L |
| GSM704528 | HSB126 | V1C | F | 30 Y | 30 | 1560 | 9.5 | R |
| GSM704866 | HSB145 | V1C | M | 36 Y | 36 | 1872 | 18 | L |
| GSM704867 | HSB145 | V1C | M | 36 Y | 36 | 1872 | 18 | R |
| GSM704461 | HSB123 | V1C | M | 37 Y | 37 | 1924 | 13 | L |
| GSM704462 | HSB123 | V1C | M | 37 Y | 37 | 1924 | 13 | R |

| | | | | | | | | |
|---|---|---|---|---|---|---|---|---|
| GSM705253 | HSB187 | V1C | F | 37 Y | 37 | 1924 | 10 | L |
| GSM705254 | HSB187 | V1C | F | 37 Y | 37 | 1924 | 10 | R |
| GSM704664 | HSB135 | V1C | F | 40 Y | 40 | 2080 | 30.5 | L |
| GSM704666 | HSB135 | V1C | F | 40 Y | 40 | 2080 | 30.5 | R |
| GSM704834 | HSB144 | V1C | M | 40 Y | 40 | 2080 | 28 | L |
| GSM704835 | HSB144 | V1C | M | 40 Y | 40 | 2080 | 28 | R |
| GSM705214 | HSB183 | V1C | M | 42 Y | 42 | 2184 | 19 | L |
| GSM705198 | HSB182 | V1C | M | 55 Y | 55 | 2860 | 13 | L |
| GSM704183 | HSB106 | V1C | M | 64 Y | 64 | 3328 | 4 | L |
| GSM704184 | HSB106 | V1C | M | 64 Y | 64 | 3328 | 4 | R |
| GSM704246 | HSB111 | V1C | F | 70 Y | 70 | 3640 | 13 | L |
| GSM704247 | HSB111 | V1C | F | 70 Y | 70 | 3640 | 13 | R |
| GSM705062 | HSB156 | V1C | F | 82 Y | 82 | 4264 | 16 | L |
| GSM705063 | HSB156 | V1C | F | 82 Y | 82 | 4264 | 16 | R |

**Supplementary Table 3**. *List of 23 proteins measured by Western blotting*. The antibodies, concentrations, suppliers, and RRIDs are listed. The citations for the publications are also included.

| Protein | Antibody Concentration and Supplier | Citation |
|---|---|---|
| GAD65 (glutamic acid decarboxylase 65) | 1:500 (Chemicon International, Temecula, CA, USA) | Pinto et al. (2010) |
| GAD67 (glutamic acid decarboxylase 67) | 1:1000 (Chemicon International, Temecula, CA, USA) | |
| VGAT | 1:1000 (Synaptic Systems, Göttingen, Germany) | |
| $GABA_A\alpha1$ | 1:500 (Imgenex, San Diego, CA, USA) | |
| $GABA_A\alpha2$ | 1:1000 (Imgenex, San Diego, CA, USA) | |
| $GABA_A\alpha3$ | 1:1000 (Imgenex, San Diego, CA, USA) | |
| CB1 | 1:1000 (Cayman, Ann Arbor, MI, USA) | |
| Gephyrin | 1:500 (Chemicon International, Temecula, CA, USA) | |
| Gephyrin Homogenate | 1:2000 (Millipore, Billerica, MA) | Pinto et al. (2015) |
| PSD95 Homogenate | 1:32000 (Millipore, Billerica, MA) | |
| Synapsin | 1:8000 (Invitrogen, Carlsbad, CA) | |
| Synaptophysin | 1:2000 (Sigma-Aldrich, St. Louis, MO) | |
| GluA2 | 1:1000 (RRID: AB_2533058, Invitrogen) | Siu et al. (2017) |
| GluN1/NR1 | 1:4000 (RRID: AB_396353, BD Biosciences PharMingen) | |
| GluN2A/NR2A | 1:1000 (RRID: AB_95169, EMD Millipore) | |
| GluN2B/NR2B | 1:1000 (RRID: AB_2112925, EMD Millipore) | |
| PSD95 | 1:16000 (RRID: AB_94278, EMD Millipore) | |
| Golli-MBP | 1:4000 [AB62631] (Abcam, Cambridge, MA, USA) | Siu et al. (2015) |
| Classic-MBP | 1:4000 [AB62631] (Abcam, Cambridge, MA, USA) | |
| Ube3A (E6AP) | 1:1000 (Bethyl Laboratories, Montgomery, TX, USA) | Williams et al. (2010) |
| Integrinβ3 | 1:1000 [AB2984] (Millipore, Billerica, MA, USA) | Unpublished |
| GFAP | 1:2000 [MAB360] (Millipore, Billerica, MA, USA) | |
| Drebrin | 1:1000 [10R-D117A] (Fitzgerald Industries International, Acton, MA, USA) | |

**Supplementary Table 4**. *List of the 95 proteins, including their gene symbol and short name*.

| Gene Symbol | Protein Name | Short Name |
|---|---|---|
| ALCAM | CD166 antigen | |
| ANG | Angiogenin | |
| AXL | Tyrosine-protein kinase receptor UFO | |
| CCL14 | C-C motif chemokine 14 | |
| CCL18 | C-C motif chemokine 18 | |
| CCL2 | C-C motif chemokine 2 | |
| CCL5 | C-C motif chemokine 5 | |
| CD14 | Monocyte differentiation antigen CD14 | |
| CEACAM1 | Carcinoembryonic antigen-related cell adhesion molecule 1 | |
| CNTN2 | Contactin-2 | |
| CSF1 | Macrophage colony-stimulating factor 1 | CSF-1 |
| CSF1R | Macrophage colony-stimulating factor 1 receptor | |
| CTSS | Cathepsin S | |
| CXCL11 | C-X-C motif chemokine 11 | |
| CXCL16 | C-X-C motif chemokine 16 | |
| CXCL8 | Interleukin-8 | IL-8 |
| EGFR | Epidermal growth factor receptor | |
| ERBB3 | Receptor tyrosine-protein kinase erbB-3 | |
| FAS | Tumor necrosis factor receptor superfamily member 6 | |
| FCGR2B | Low affinity immunoglobulin gamma Fc region receptor II-b | IgG Fc receptor II-b |
| FGF2 | Fibroblast growth factor 2 | FGF-2 |
| FLT1 | Vascular endothelial growth factor receptor 1 | VEGFR-1 |
| FLT3LG | Fms-related tyrosine kinase 3 ligand | Flt3 ligand |
| GDF15 | Growth/differentiation factor 15 | GDF-15 |
| GDNF | Glial cell line-derived neurotrophic factor | hGDNF |
| HGF | Hepatocyte growth factor | |
| ICAM1 | Intercellular adhesion molecule 1 | ICAM-1 |
| ICAM2 | Intercellular adhesion molecule 2 | ICAM-2 |
| IGFBP1 | Insulin-like growth factor-binding protein 1 | IBP-1 |
| IGFBP2 | Insulin-like growth factor-binding protein 2 | IBP-2 |
| IGFBP6 | Insulin-like growth factor-binding protein 6 | IBP-6 |
| IL11 | Interleukin-11 | IL-11 |
| IL13RA1 | Interleukin-13 receptor subunit alpha-1 | IL-13 receptor subunit alpha-1 |

| | | |
|---|---|---|
| IL13RA2 | Interleukin-13 receptor subunit alpha-2 | IL-13 receptor subunit alpha-2 |
| IL16 | Pro-interleukin-16 | |
| IL1A | Interleukin-1 alpha | IL-1 alpha |
| IL1RN | Interleukin-1 receptor antagonist protein | IL-1RN |
| IL2RB | Interleukin-2 receptor subunit beta | IL-2 receptor subunit beta |
| IL2RG | Cytokine receptor common subunit gamma | |
| IL4 | Interleukin-4 | IL-4 |
| IL6R | Interleukin-6 receptor subunit alpha | IL-6 receptor subunit alpha |
| IL6ST | Interleukin-6 receptor subunit beta | IL-6 receptor subunit beta |
| IL9 | Interleukin-9 | IL-9 |
| KIT | Mast/stem cell growth factor receptor Kit | SCFR |
| LCN2 | Neutrophil gelatinase-associated lipocalin | NGAL |
| LIF | Leukemia inhibitory factor | LIF |
| LYVE1 | Lymphatic vessel endothelial hyaluronic acid receptor 1 | LYVE-1 |
| MICA | MHC class I polypeptide-related sequence A | MIC-A |
| NRCAM | Neuronal cell adhesion molecule | Nr-CAM |
| PDGFA | Platelet-derived growth factor subunit A | PDGF subunit A |
| PDGFAB | Platelet-derived growth factor subunit AB | PDGF subunit AB |
| PDGFB | Platelet-derived growth factor subunit B | PDGF subunit B |
| PECAM1 | Platelet endothelial cell adhesion molecule | PECAM-1 |
| PF4 | Platelet factor 4 | PF-4 |
| PI3 | Elafin | |
| PLAUR | Urokinase plasminogen activator surface receptor | U-PAR |
| PPBP | Platelet basic protein | PBP |
| PROK1 | Prokineticin-1 | |
| RETN | Resistin | |
| SCARB2 | Lysosome membrane protein 2 | |
| SIGLEC5 | Sialic acid-binding Ig-like lectin 5 | Siglec-5 |
| SPP1 | Osteopontin | |
| TGFB1 | Human TGF-beta 1 cDNA | |
| TGFA | Protransforming growth factor alpha | |
| TIMP1 | Metalloproteinase inhibitor 1 | |
| TIMP2 | Metalloproteinase inhibitor 2 | |
| TNF | Tumor necrosis factor | |
| TNFRSF10C | Tumor necrosis factor receptor superfamily member 10C | |

| | | |
|---|---|---|
| TNFRSF14 | Tumor necrosis factor receptor superfamily member 14 | |
| TNFRSF1A | Tumor necrosis factor receptor superfamily member 1A | |
| TNFRSF1B | Tumor necrosis factor receptor superfamily member 1B | |
| TYRO3 | Tyrosine-protein kinase receptor TYRO3 | |
| CNR1 | Cannabinoid receptor 1 | CB-R |
| MBP | Classic myelin basic protein | MBP |
| DBN1 | Drebrin | |
| GABRA1 | Gamma-aminobutyric acid receptor subunit alpha-1 | |
| GABRA2 | Gamma-aminobutyric acid receptor subunit alpha-2 | |
| GABRA3 | Gamma-aminobutyric acid receptor subunit alpha-3 | |
| GAD2 | Glutamate decarboxylase 2 | |
| GAD1 | Glutamate decarboxylase 1 | |
| GPHN | Gephyrin | |
| GPHN | Gephyrin (HOM) | |
| GFAP | Glial fibrillary acidic protein | GFAP |
| GRIA2 | Glutamate receptor 2 | GluR-2 |
| GRIN1 | Glutamate receptor ionotropic, NMDA 1 | GluN1 |
| MBP | Golli myelin basic protein | Golli MBP |
| ITGB3 | Integrin beta-3 | |
| GRIN2A | Glutamate receptor ionotropic, NMDA 2A | GluN2A |
| GRIN2B | Glutamate receptor ionotropic, NMDA 2B | GluN2B |
| DLG4 | Disks large homolog 4 | PSD-95 |
| DLG4 | Disks large homolog 4 (HOM) | PSD-95 |
| SYN1 | Synapsin-1 | NA |
| SYP | Synaptophysin | NA |
| UBE3A | Ubiquitin-protein ligase E3A | NA |
| SLC32A1 | Vesicular inhibitory amino acid transporter | hVIAAT |

Supplementary Figure S1. Clusters when the age of the samples are randomized.

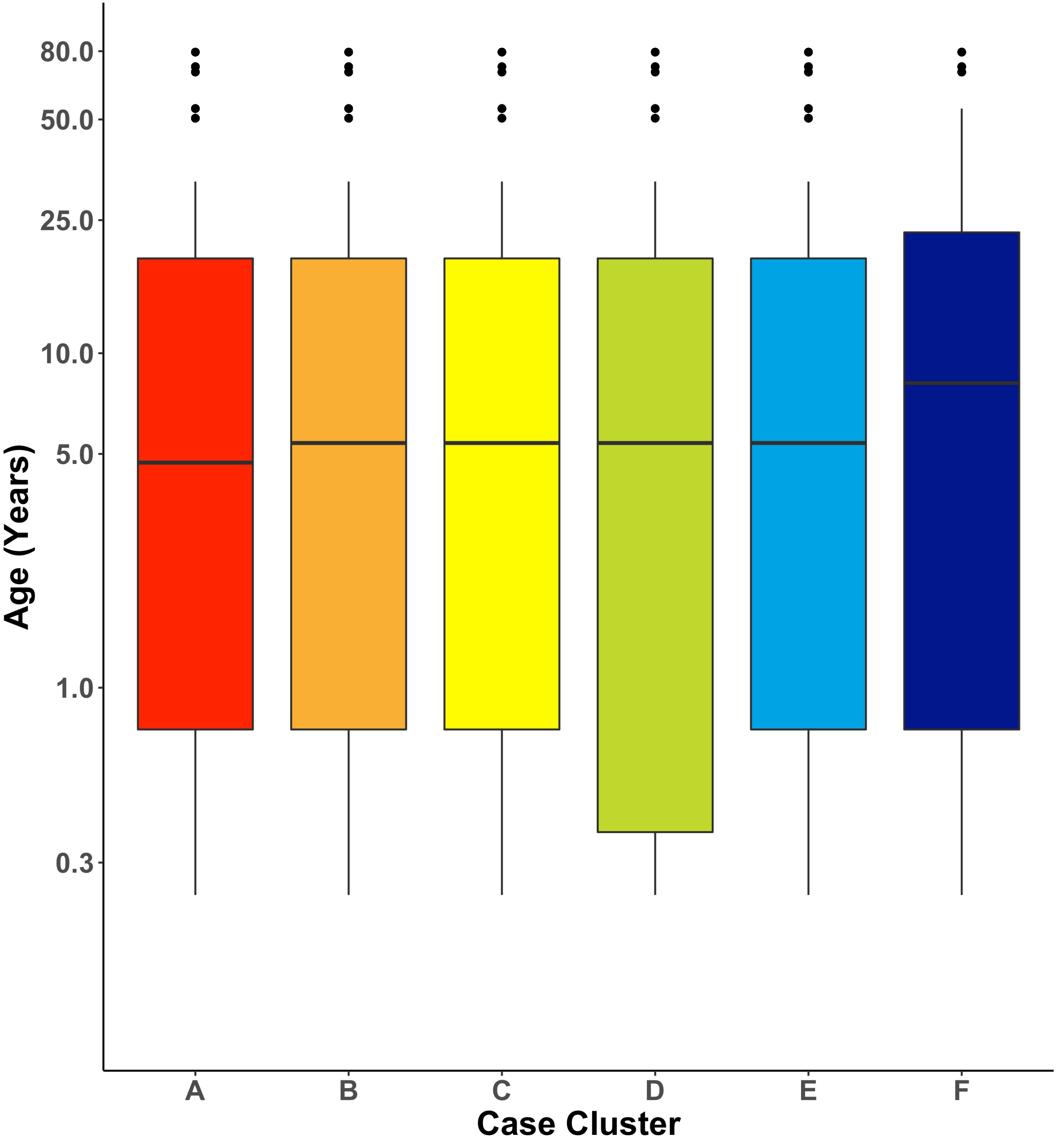

Supplementary Figure S2. Normalized expression and age binned for the 88 gene and protein pairs.

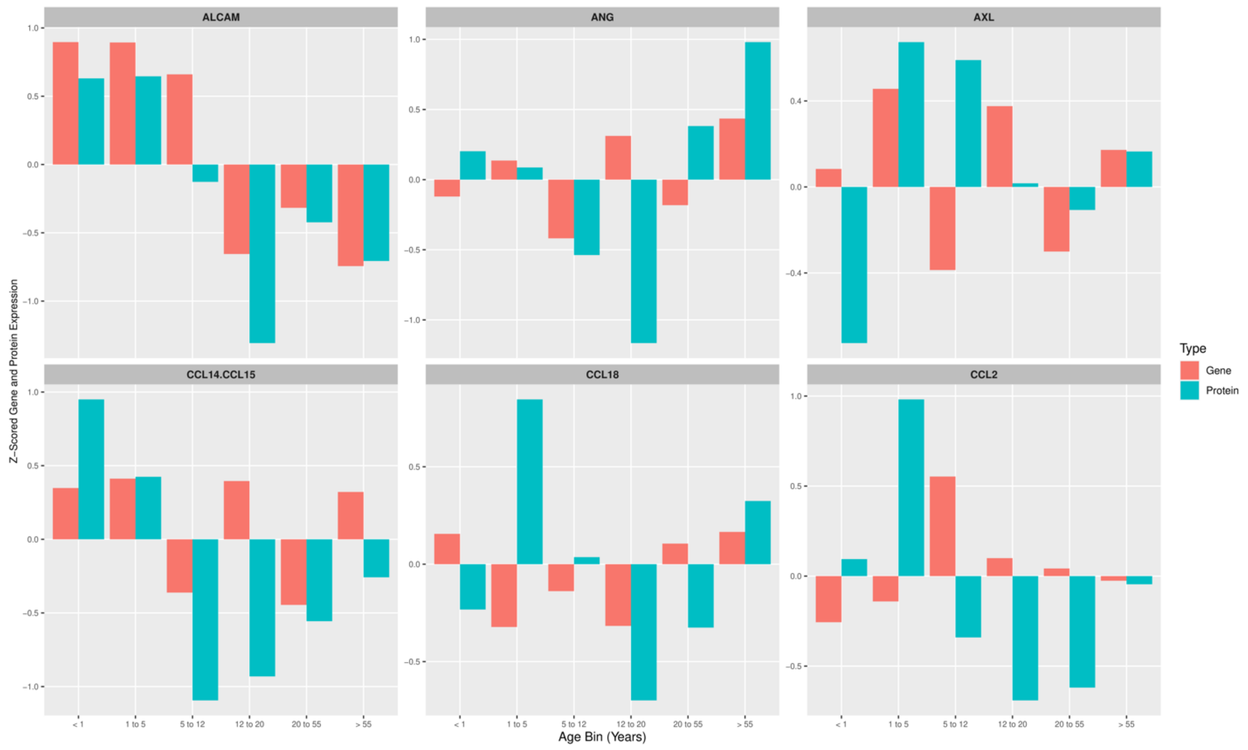


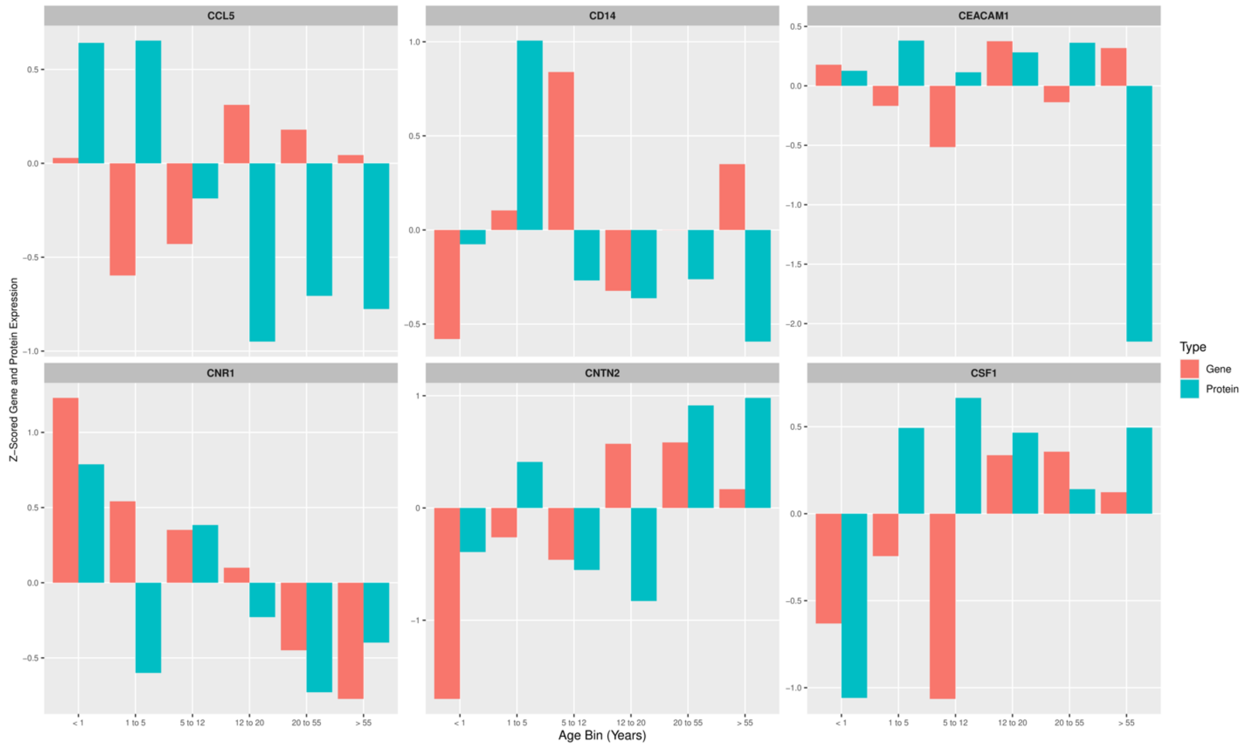

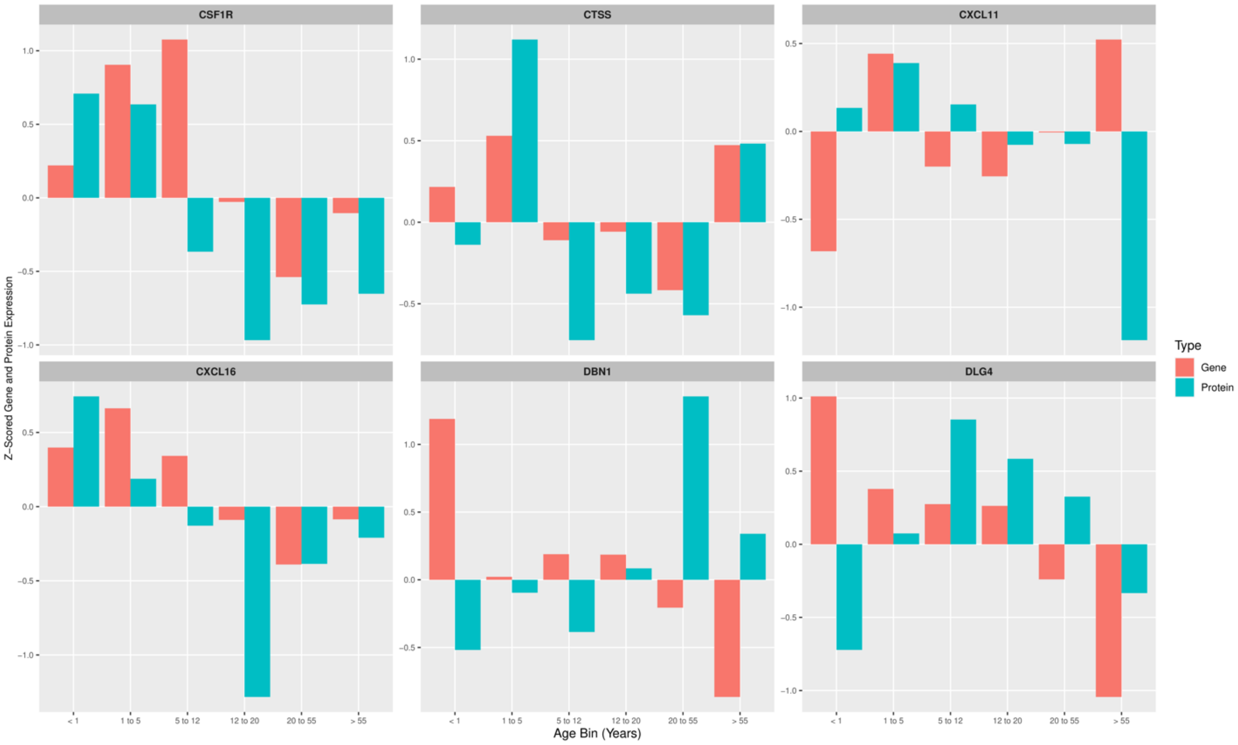
CSF1R
CTSS
CXCL11
CXCL16
DBN1
DLG4
Z-Scored Gene and Protein Expression
< 1
1 to 5
5 to 12
12 to 20
20 to 55
> 55
Age Bin (Years)
Type
Gene
Protein

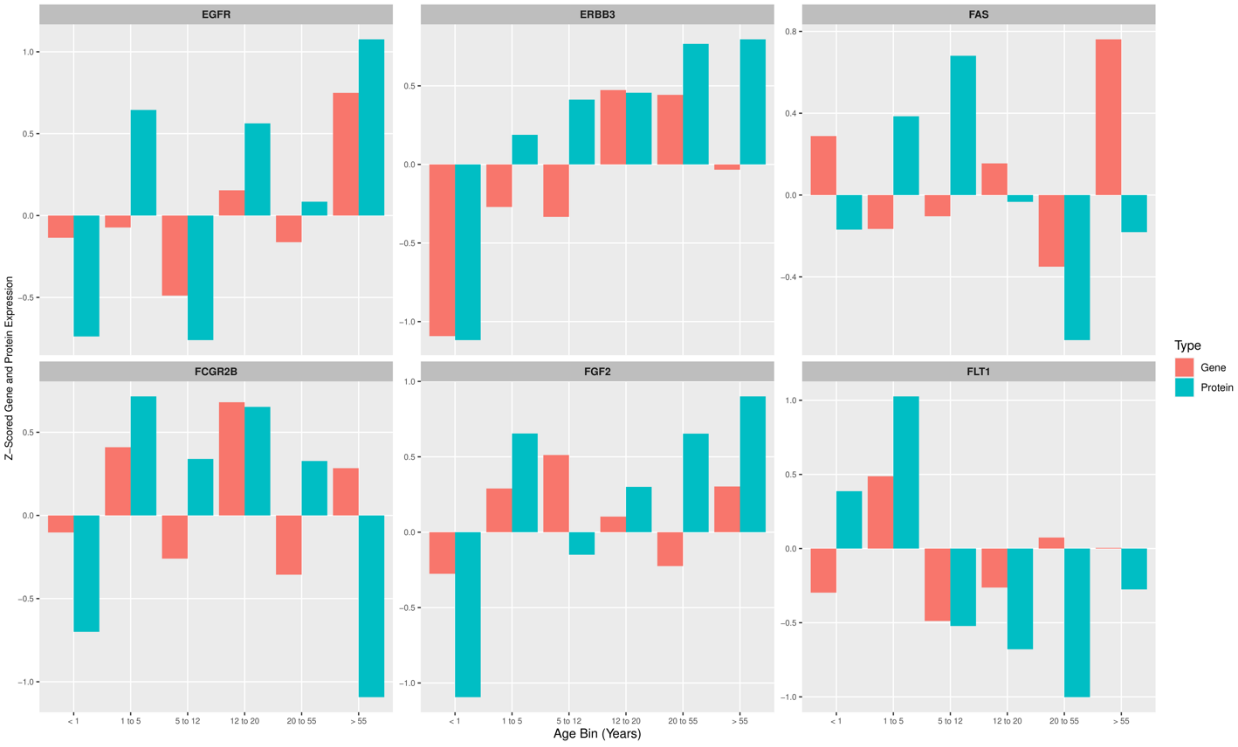
EGFR
ERBB3
FAS
FCGR2B
FGF2
FLT1
Z-Scored Gene and Protein Expression
< 1
1 to 5
5 to 12
12 to 20
20 to 55
> 55
Age Bin (Years)
Type
Gene
Protein

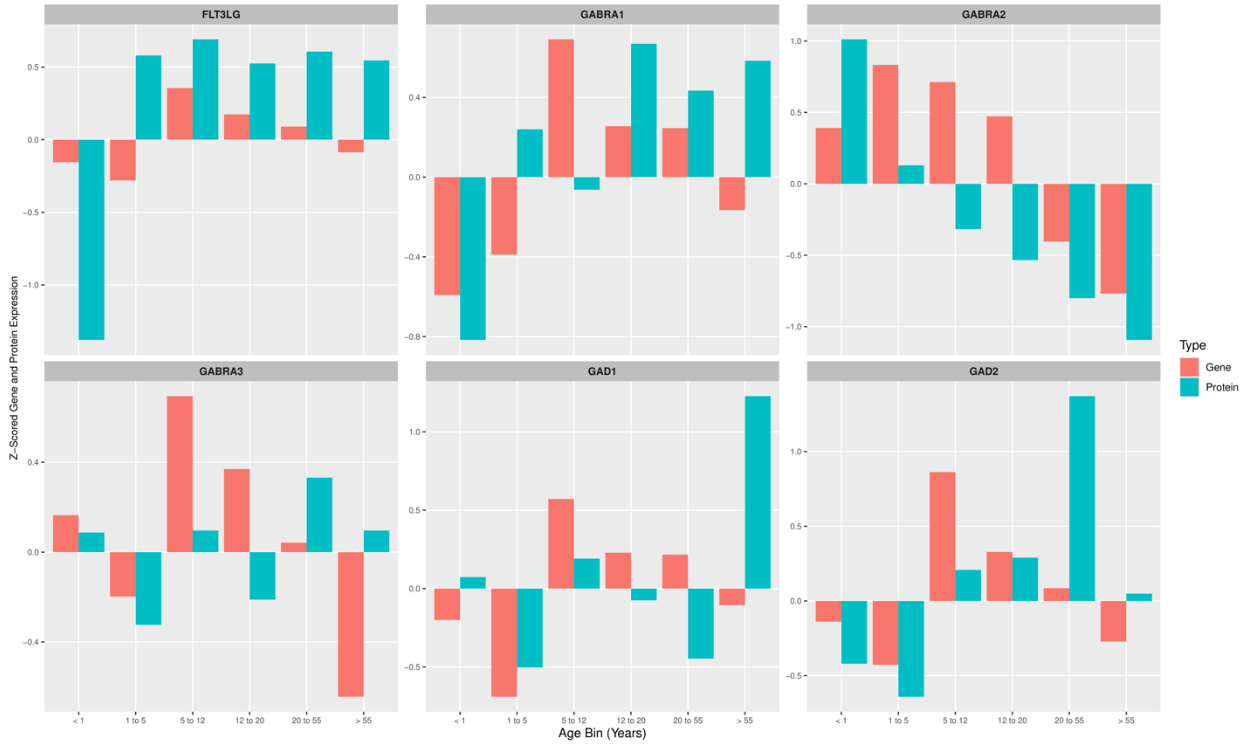
FLT3LG
GABRA1
GABRA2
GABRA3
GAD1
GAD2
Z-Scored Gene and Protein Expression
Age Bin (Years)
< 1
1 to 5
5 to 12
12 to 20
20 to 55
> 55
Type
Gene
Protein

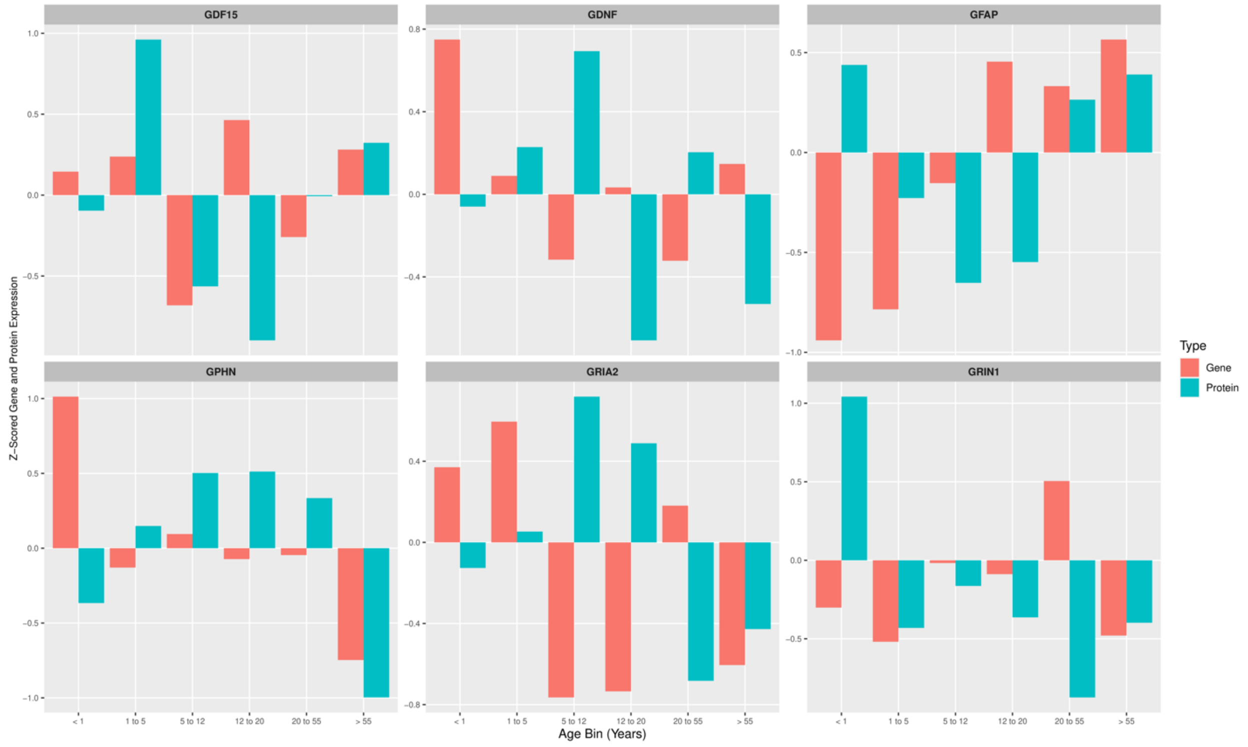
GDF15
GDNF
GFAP
GPHN
GRIA2
GRIN1
Z-Scored Gene and Protein Expression
Age Bin (Years)
< 1
1 to 5
5 to 12
12 to 20
20 to 55
> 55
Type
Gene
Protein

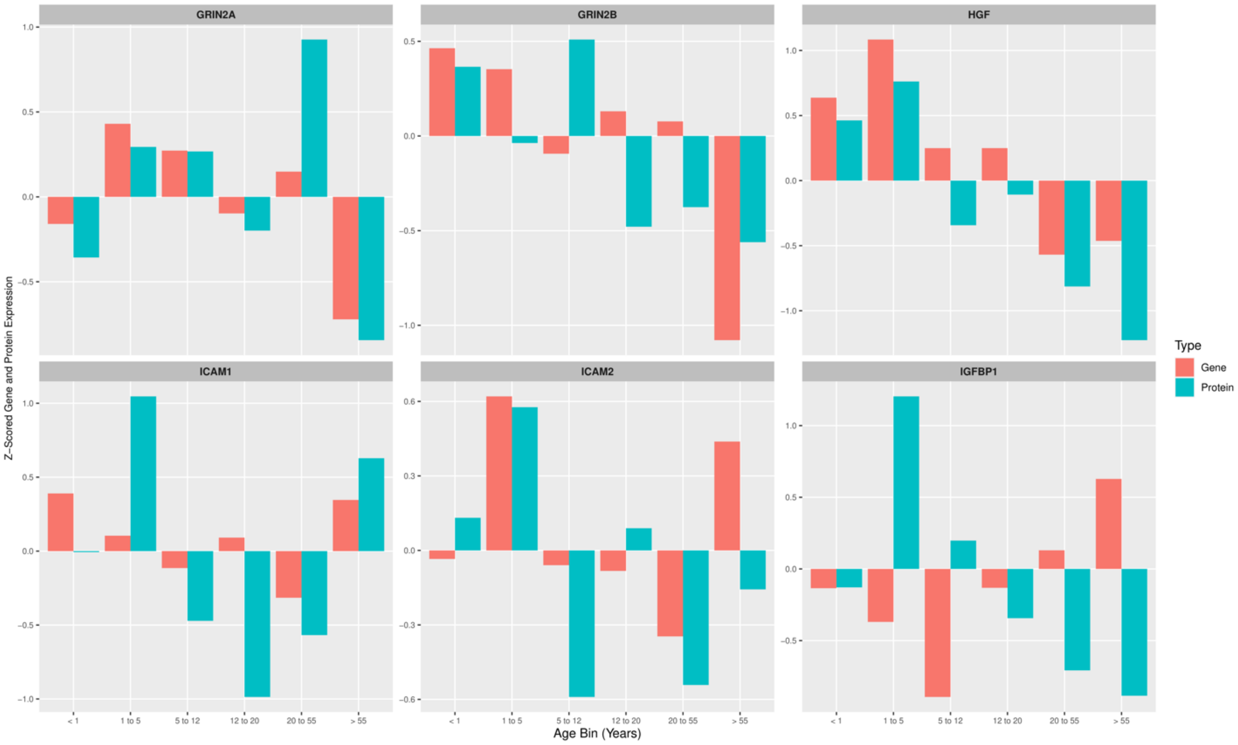
GRIN2A
GRIN2B
HGF
ICAM1
ICAM2
IGFBP1
Z−Scored Gene and Protein Expression
Age Bin (Years)
< 1
1 to 5
5 to 12
12 to 20
20 to 55
> 55
Type
Gene
Protein

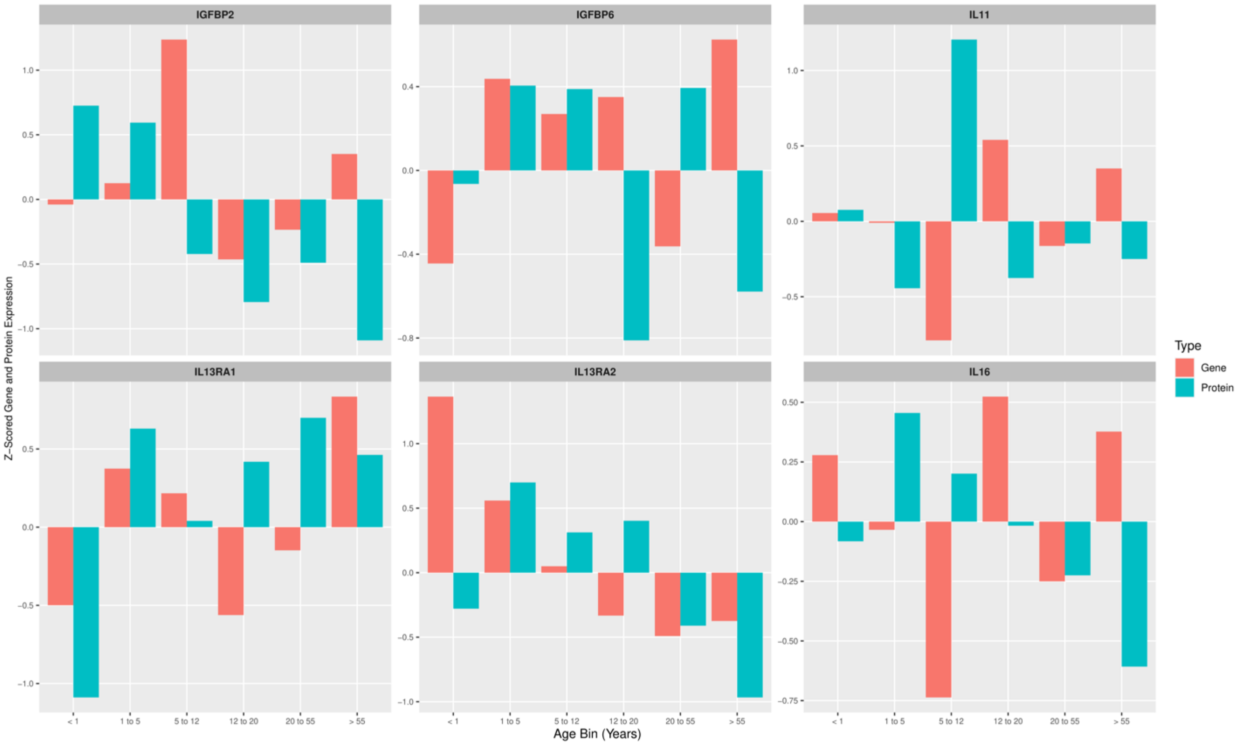
IGFBP2
IGFBP6
IL11
IL13RA1
IL13RA2
IL16
Z−Scored Gene and Protein Expression
Age Bin (Years)
< 1
1 to 5
5 to 12
12 to 20
20 to 55
> 55
Type
Gene
Protein

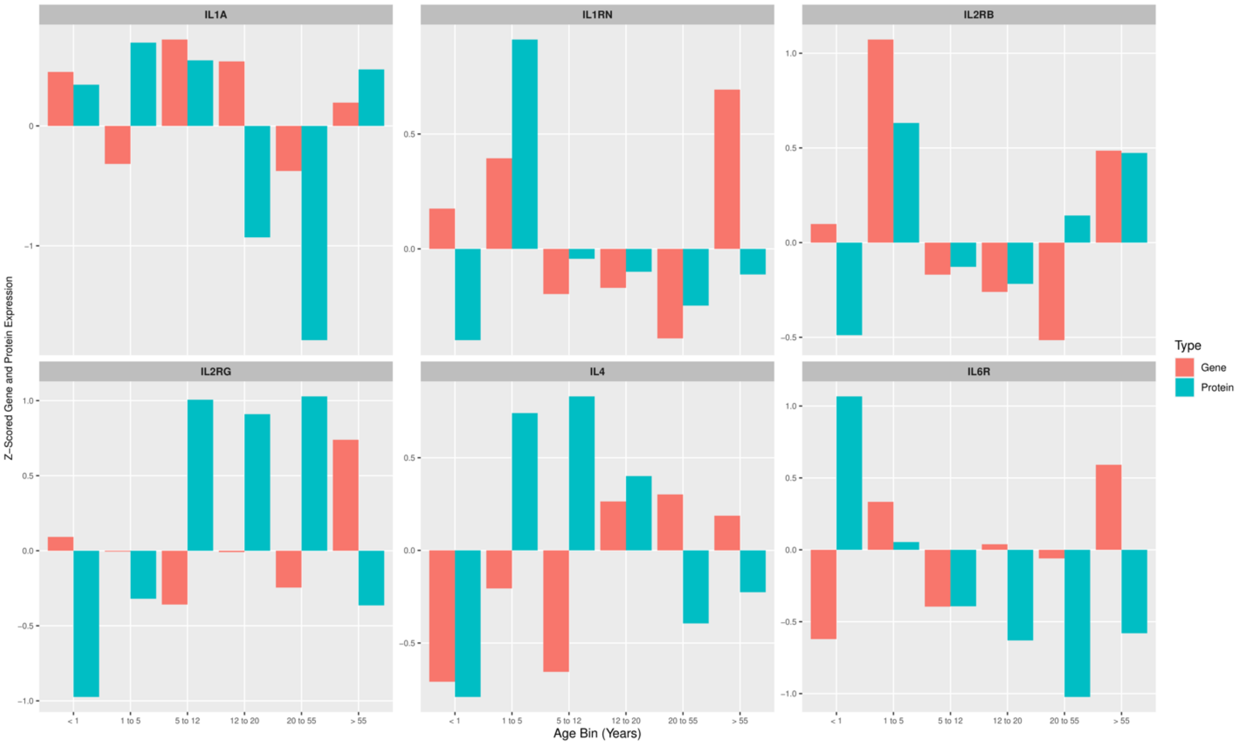
IL1A
IL1RN
IL2RB
IL2RG
IL4
IL6R
Z−Scored Gene and Protein Expression
Age Bin (Years)
< 1
1 to 5
5 to 12
12 to 20
20 to 55
> 55
Type
Gene
Protein

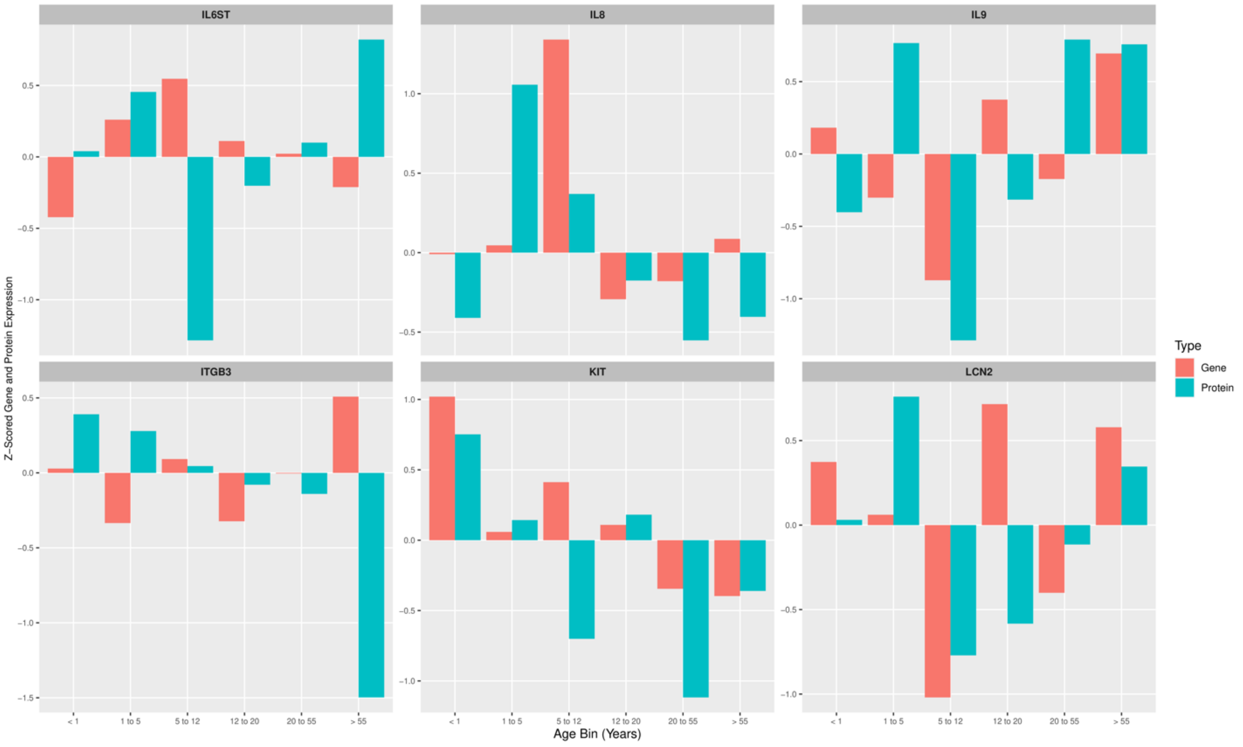
IL6ST
IL8
IL9
ITGB3
KIT
LCN2
Z−Scored Gene and Protein Expression
Age Bin (Years)
< 1
1 to 5
5 to 12
12 to 20
20 to 55
> 55
Type
Gene
Protein

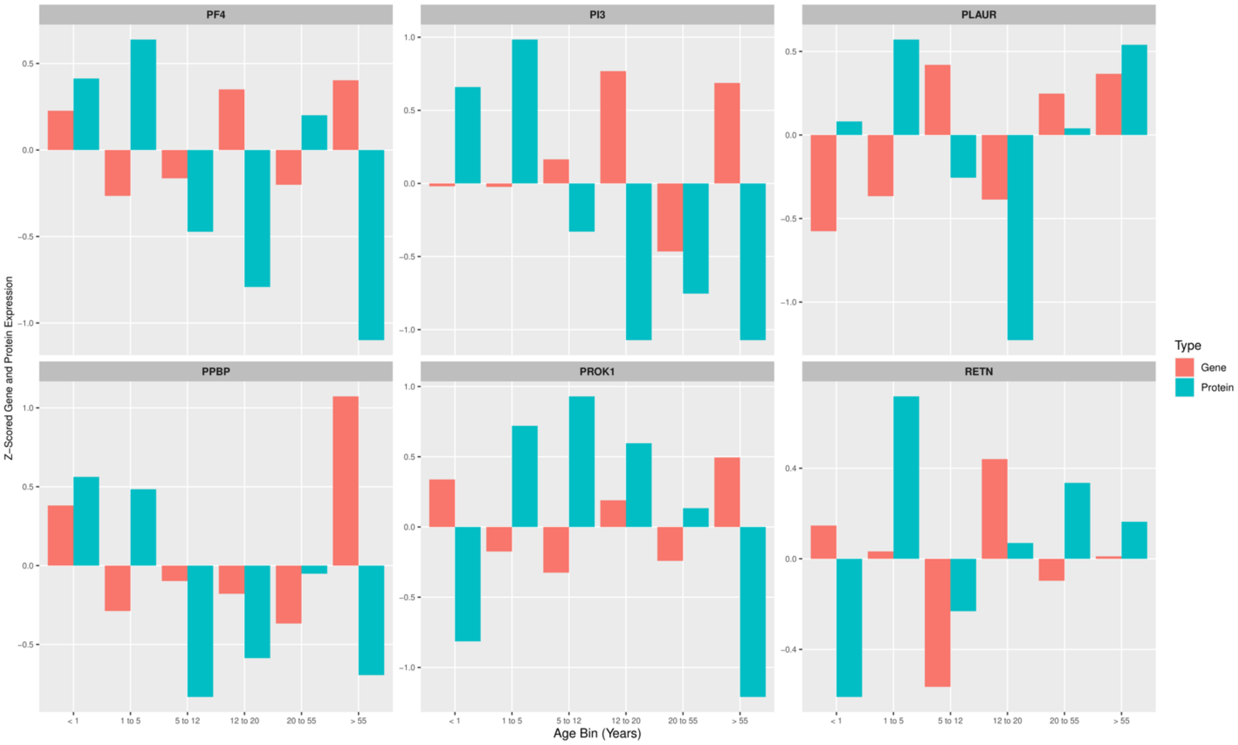
PF4
PI3
PLAUR
PPBP
PROK1
RETN
Z-Scored Gene and Protein Expression
< 1
1 to 5
5 to 12
12 to 20
20 to 55
> 55
Age Bin (Years)
Type
Gene
Protein

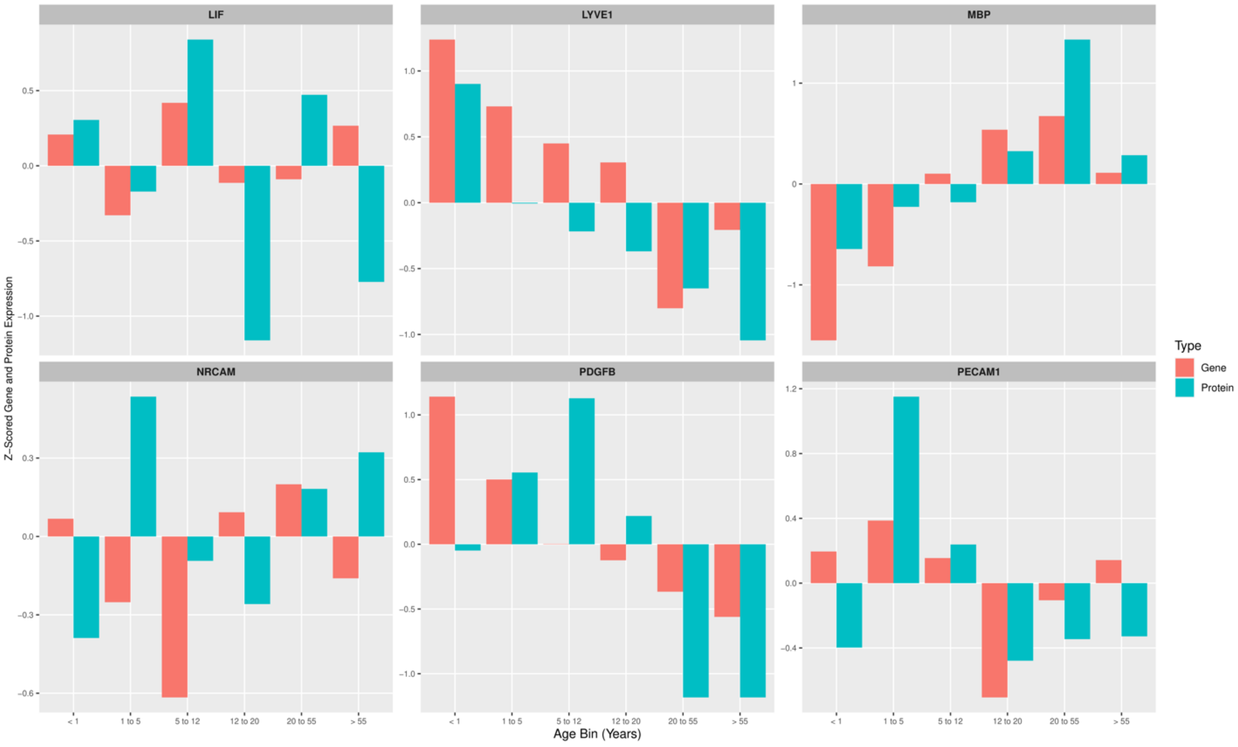
LIF
LYVE1
MBP
NRCAM
PDGFB
PECAM1
Z-Scored Gene and Protein Expression
< 1
1 to 5
5 to 12
12 to 20
20 to 55
> 55
Age Bin (Years)
Type
Gene
Protein

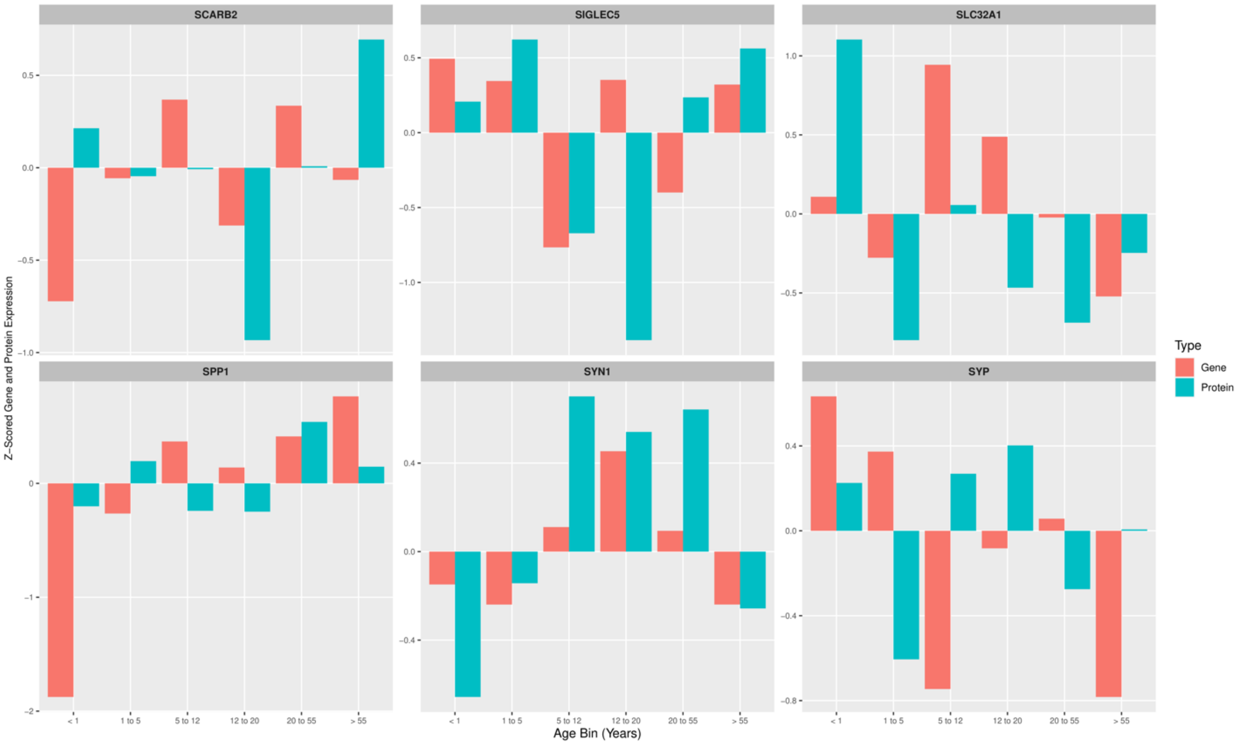
SCARB2
SIGLEC5
SLC32A1
SPP1
SYN1
SYP
Z-Scored Gene and Protein Expression
< 1
1 to 5
5 to 12
12 to 20
20 to 55
> 55
Age Bin (Years)
Type
Gene
Protein

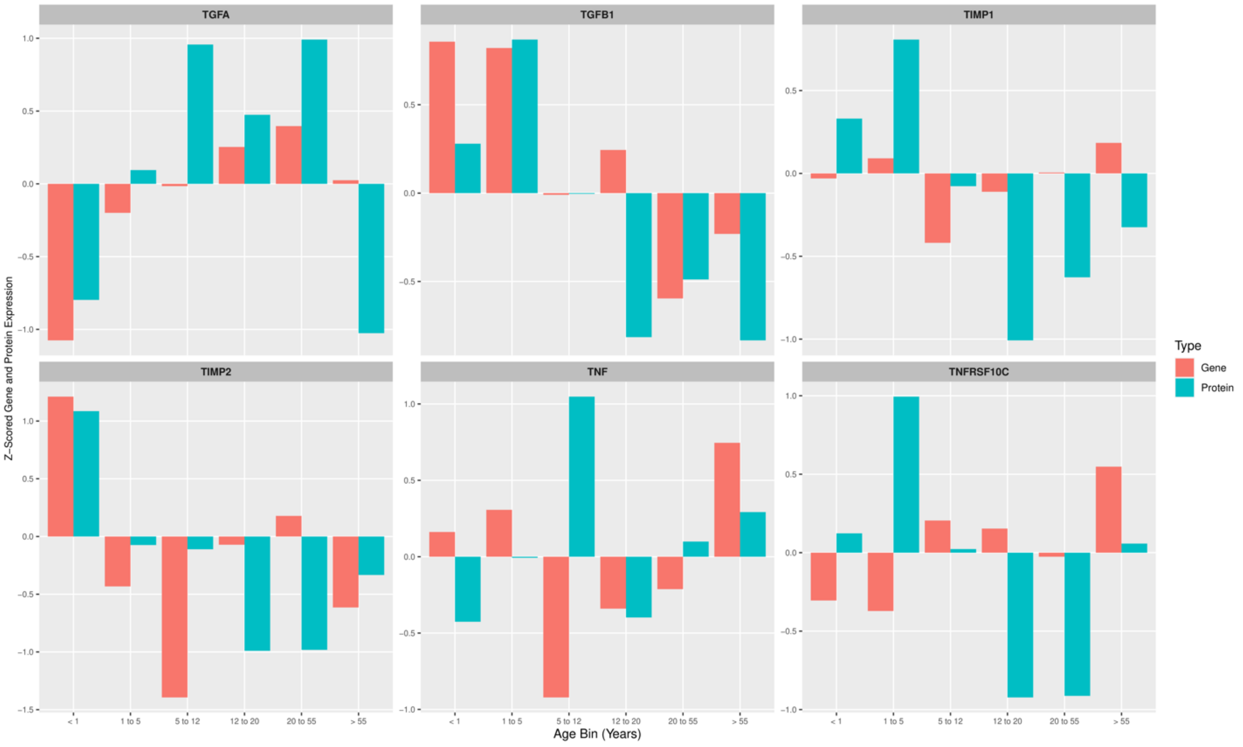
TGFA
TGFB1
TIMP1
TIMP2
TNF
TNFRSF10C
Z-Scored Gene and Protein Expression
< 1
1 to 5
5 to 12
12 to 20
20 to 55
> 55
Age Bin (Years)
Type
Gene
Protein

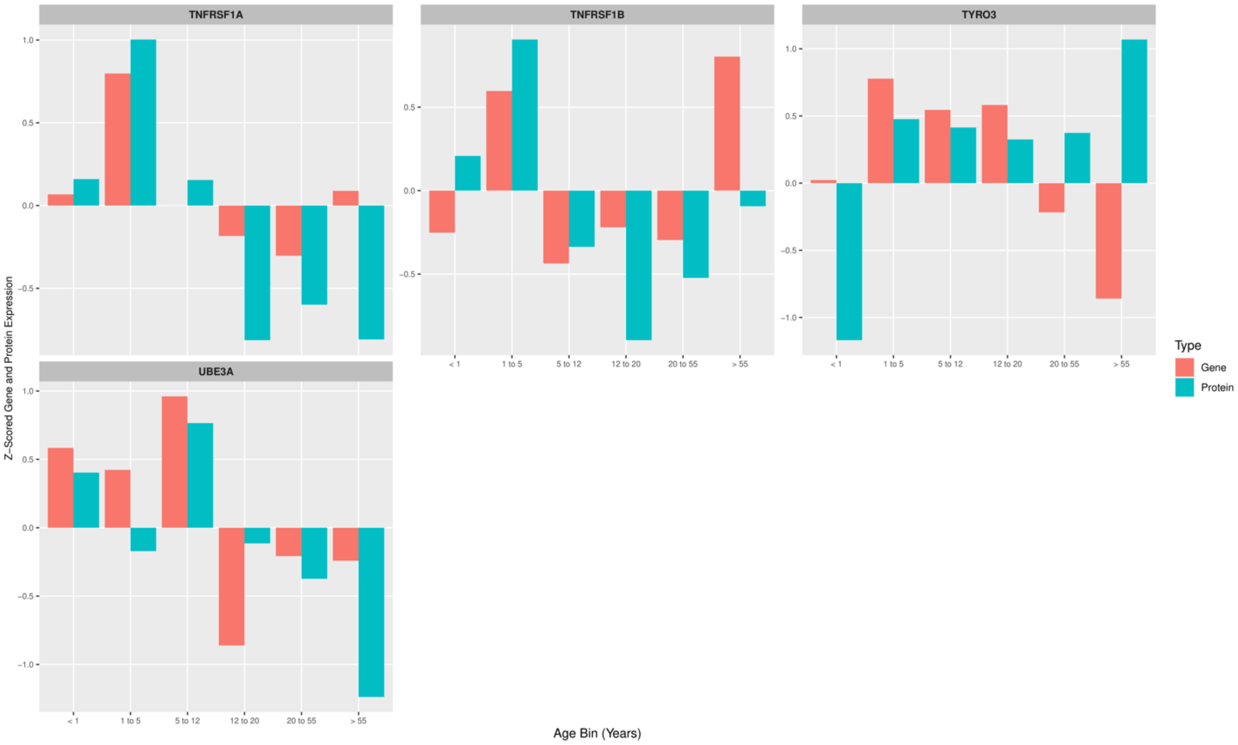


**Supplementary Figure S2**. *Bar plots of normalized expression for 88 gene and protein pairs across age bins*. Individual facet plots are labelled by the corresponding gene. Gene expression is represented by red bars and protein expression is shown with blue bars.

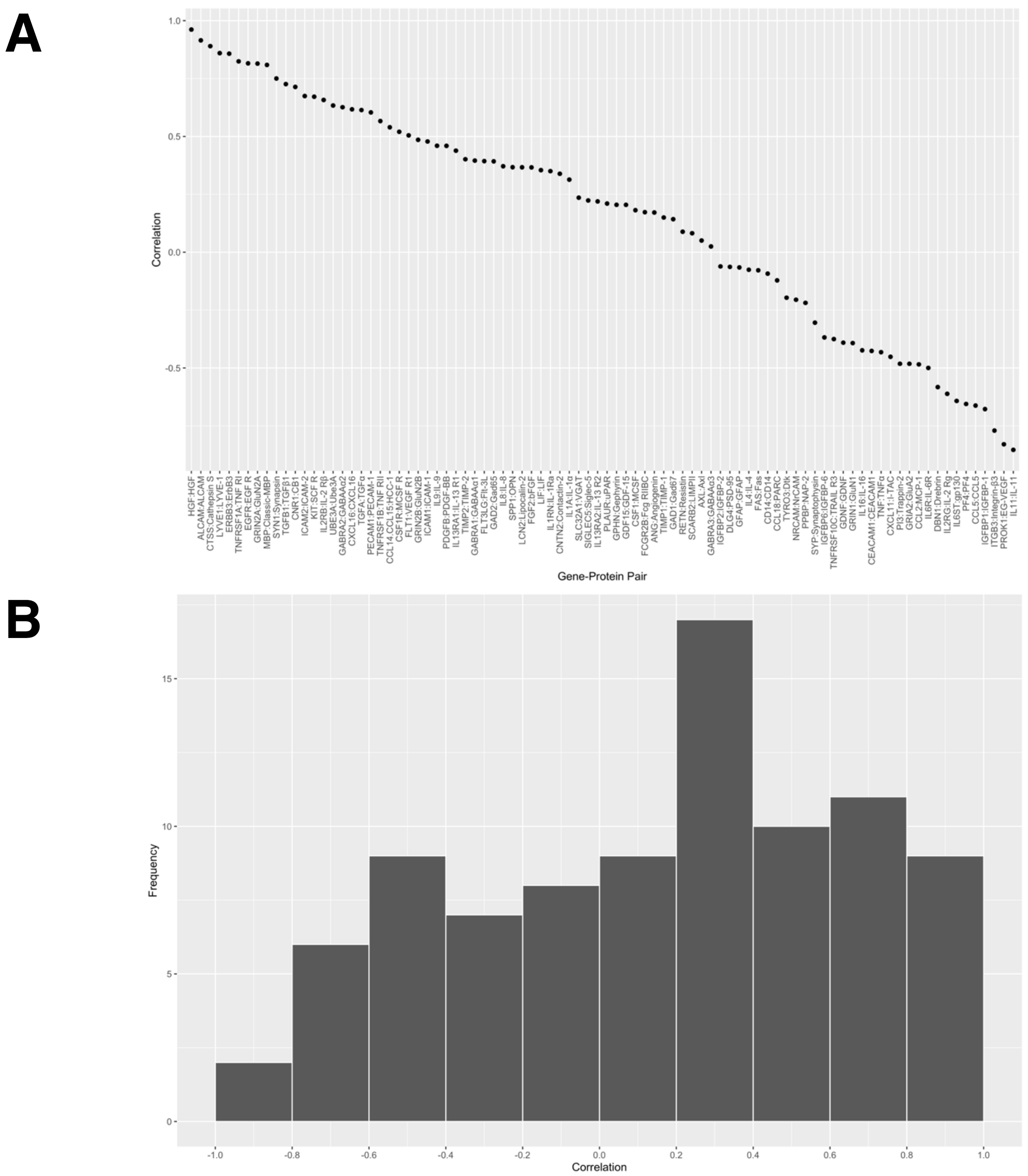


**Supplementary Figure S3**. **A)** *Scatterplot of the correlations obtained between corresponding gene and protein pairs*. Points represent Pearson's R correlation coefficients calculated for the expression of each of 88 gene-protein pairs, arranged in decreasing order. **B)** *Histogram of correlations obtained from expression of 88 gene-protein pairs*. Bins for the correlation are shown in increments of 0.2.

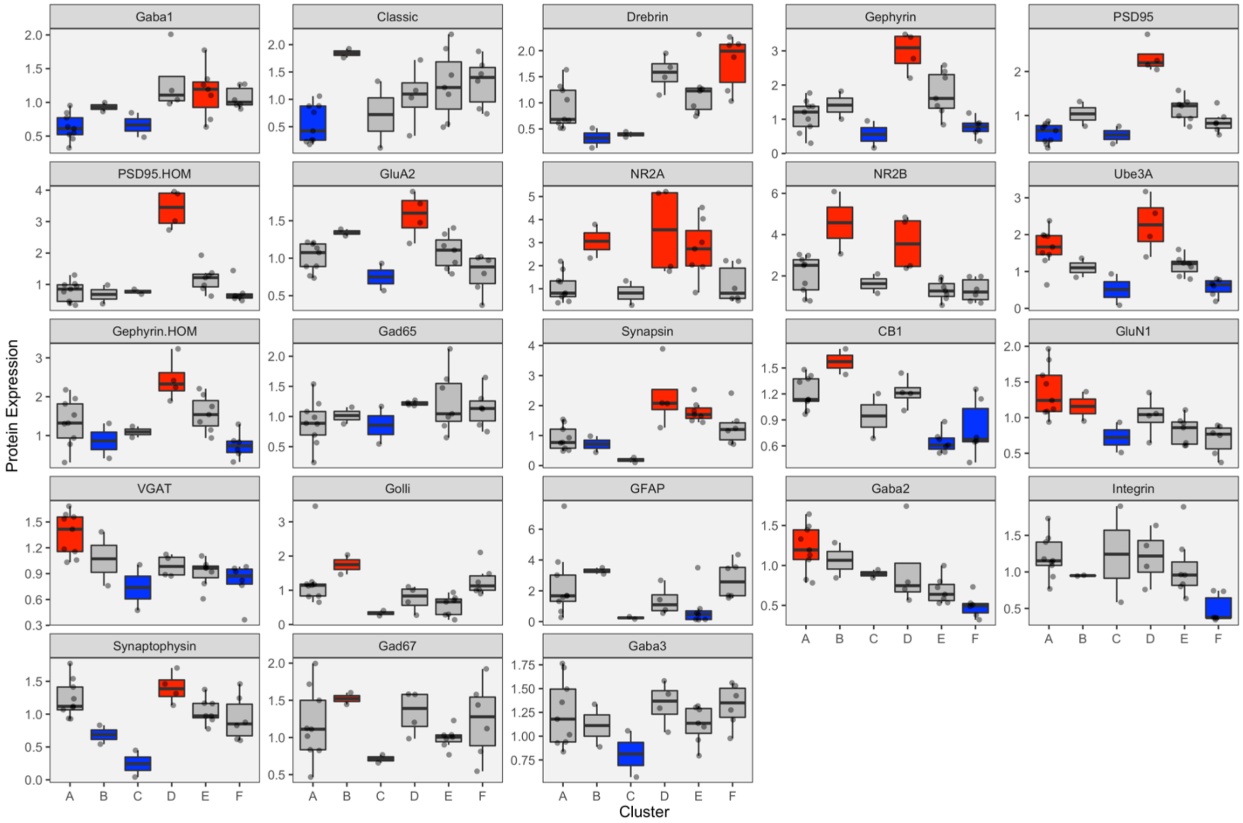


**Supplementary Figure S4**. *Boxplots show the expression of each protein within each cluster, with an alternate rule for identifying over- or under-representation*. Red coloured boxes represent the median expression of a group being greater than the upper quantile (i.e. 75$^{th}$ percentile) of all the data. Blue coloured boxes represent the median expression of a group being less than the bottom quantile (i.e. 25$^{th}$ percentile) of all the data. Boxes are grey if their median expression value is between the lower and upper quantiles.

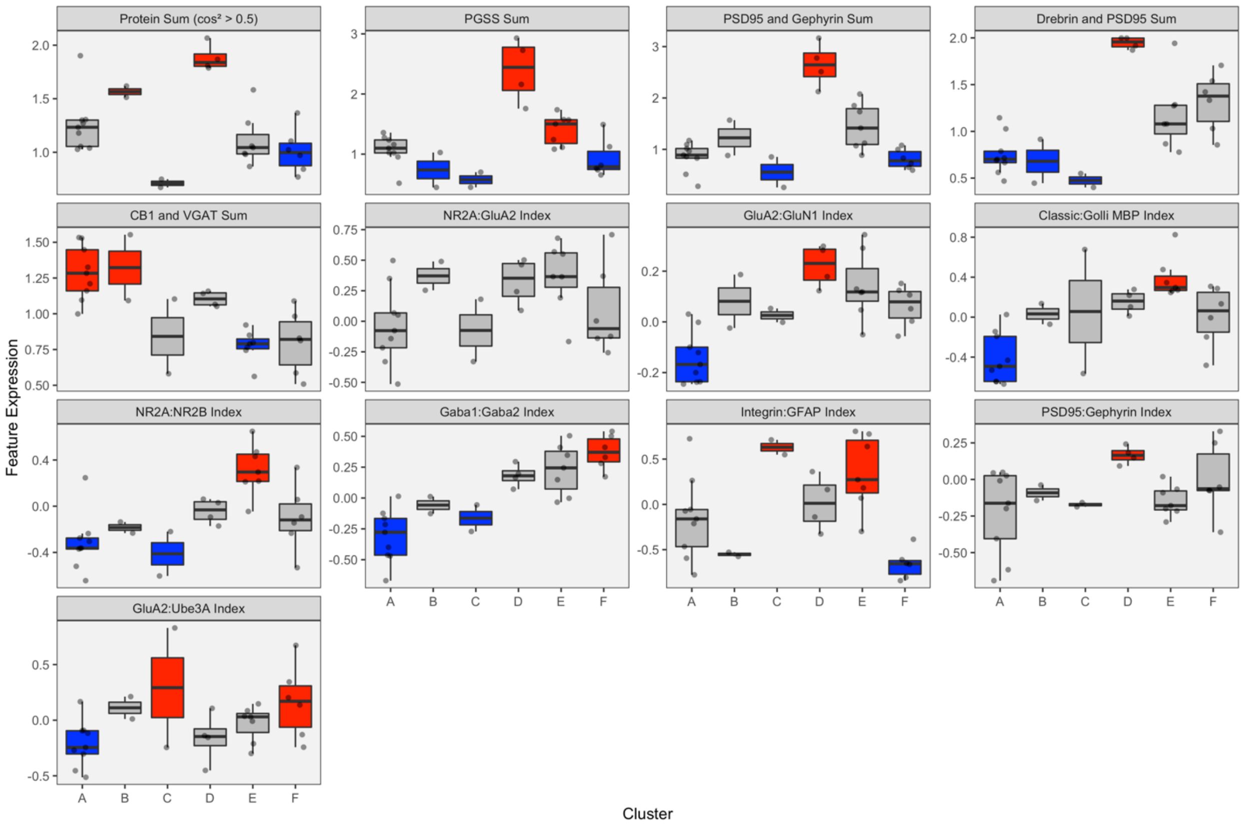


**Supplementary Figure S5**. *Boxplots show the expression of each feature within each cluster, with an alternate rule for identifying over- or under-representation*. Red coloured boxes represent the median expression of a group being greater than the upper quantile (i.e. 75th percentile) of all the data. Blue coloured boxes represent the median expression of a group being less than the bottom quantile (i.e. 25th percentile) of all the data. Boxes are grey if their median expression value is between the lower and upper quantiles.
